# Glycomic analysis of host-response reveals high mannose as a key mediator of influenza severity

**DOI:** 10.1101/2020.04.21.054098

**Authors:** Daniel W. Heindel, Sujeethraj Koppolu, Yue Zhang, Brian Kasper, Lawrence Meche, Christopher Vaiana, Stephanie J. Bissel, Chalise E. Carter, Alyson A. Kelvin, Bin Zhang, Bin Zhou, Tsui-Wen Chou, Lauren Lashua, Ted M. Ross, Elodie Ghedin, Lara K. Mahal

## Abstract

Influenza virus infections cause a wide variety of outcomes, from mild disease to 3-5 million cases of severe illness and ~290,000-645,000 deaths annually worldwide. The molecular mechanisms underlying these disparate outcomes are currently unknown. Glycosylation within the human host plays a critical role in influenza virus biology. However, the impact these modifications have on the severity of influenza disease has not been examined. Herein, we profile the glycomic host responses to influenza virus infection as a function of disease severity using a ferret model and our lectin microarray technology. We identify the glycan epitope high mannose as a marker of influenza virus-induced pathogenesis and severity of disease outcome. Induction of high mannose is dependent upon the unfolded protein response (UPR) pathway, a pathway previously shown to associate with lung damage and severity of influenza virus infection. Also, the mannan-binding lectin (MBL2), an innate immune lectin that negatively impacts influenza outcomes, recognizes influenza virus-infected cells in a high mannose dependent manner. Together, our data argue that the high mannose motif is an infection-associated molecular pattern on host cells that may guide immune responses leading to the concomitant damage associated with severity.

**SIGNIFICANCE:** Influenza virus infection causes a range of outcomes from mild illness to death. The molecular mechanisms leading to these differential host responses are currently unknown. Herein, we identify the induction of high mannose, a glycan epitope, as a key mediator of severe disease outcome. We propose a mechanism in which activation of the unfolded protein response (UPR) upon influenza virus infection turns on expression of high mannose, which is then recognized by the innate immune lectin MBL2, activating the complement cascade and leading to subsequent inflammation. This work is the first to systematically study host glycomic changes in response to influenza virus infection, identifying high mannose as a key feature of differential host response.

## INTRODUCTION

Influenza virus infections cause ~290,000-645,000 deaths (1) and 3-5 million cases of severe illness annually worldwide (2). Viral infection begins in the upper respiratory tract. It can lead to airway inflammation in the lungs and, in severe cases, pneumonia. The molecular mechanisms that underlie disparate outcomes of mild, moderate, and severe illness are currently unknown.

Glycosylation within the human host plays a critical role in influenza biology (3). Influenza virus infection is initiated by adhesion of viral hemagglutinin (HA) to glycans containing α-2,6-linked sialic acids on the human host epithelial cells. Viral propagation requires cleavage of these sialic acid residues by the influenza virus enzyme neuraminidase (NA), the current target of some anti-viral therapies (4). Due to their role in influenza biology, glycans have been studied in influenza almost exclusively in the context of binding to sialic acids. However, whether sialic acid plays a role in the severity of the host response to influenza virus infection is unknown. In addition, there are currently no systematic studies on the response of the host glycome to influenza virus infection. Whether host glycosylation (e.g. sialylation and other motifs) is modulated upon infection, and what impact these modifications could have on the severity of influenza virus induced disease have not been examined.

Studies of influenza in model organisms, such as mice, commonly use strains of the virus adapted to the sialic acid receptors prevalent in the upper respiratory tract of these hosts (i.e. α-2,3-linked sialosides), which differ from α-2,6-sialosides found in the human respiratory tract. Ferrets, however, have a similar glycan distribution in their respiratory system to humans, *i.e.* α-2,6-linked sialic acids and low levels of N-glycolyl sialosides in the upper respiratory tract. This makes them a more representative model for human biology, enabling the use of un-adapted human strains of influenza (5).

Here, we ask whether glycosylation might play a role in the outcome of influenza virus infection. We analyzed ferret host response in the lung to the 2009 pandemic H1N1 (H1N1pdm09) influenza virus using our lectin microarray technology (6–8). This H1N1 strain caused an estimated 150,000-580,000 deaths worldwide in the first 12 months of circulation, with wide variation in outcomes among infected individuals (9). Our lectin microarray technology is a well-established method that provides a systems-level perspective on glycosylation (6, 8, 10, 11). Although we observed changes in sialic acid in response to H1N1pdm09 influenza virus infection, they did not correlate with severity. Instead, we identified high mannose, an epitope rarely observed at the cell surface (12), as a marker of severity and damage in the ferret lung. Induction of high mannose was shown to depend upon the unfolded protein response (UPR) pathway, which is associated with influenza disease severity in mouse models (13). We also observed that mannan-binding lectin (MBL2), an innate immune lectin that binds several different glycans (*e.g.* Lewis structures (14, 15), high mannose (16), yeast mannans (17), fucose (15)), recognizes influenza virus infected cells in a high mannose dependent manner. MBL2 is also associated with the severity of influenza in mouse models (18) and more recently in clinical analysis (19). Together, our data argue that the high mannose motif is an influenza infection-associated molecular pattern on host cells that guides immune responses leading to the concomitant damage associated with influenza disease severity.

## RESULTS

### Lectin Microarray Analysis of Ferret Lungs After Influenza Infection

To study the impact of influenza virus infection on host glycosylation, we infected ferrets with the 2009 pandemic H1N1 strain, A/California/07/2009 (H1N1). This strain causes a wide variation in outcomes among infected ferrets, mimicking the human host response (20). Infected adult ferrets (n=19) were weighed daily during the experiment and sacrificed at day 8 post-infection. By day 8, ferrets are at the start of the recovery period from infection, as observed from weight loss and pathology data (20). The severity of the infection was determined based on weight loss. The weight loss nadir was used with the lowest quartile defined as mild (weight loss less than 10.5%, n=5 ferrets), the middle quartile as moderate (weight loss between 10.5% and 16.2%, n=8 ferrets) and the highest quartile as severe (weight loss greater than 16.2%, n=6 ferrets).

We analyzed lung punch biopsies from both the upper and lower lungs of infected animals at day 8 post-infection (n=2 samples per animal) using our dual-color lectin microarray method. Lectin microarrays use carbohydrate-binding proteins of known specificities as probes for glycan structure and provide a systems-level view of the glycome (6–8). For control animals, additional biopsy locations (n=6 samples total per animal, 4 ferrets) were analyzed for additional statistical power. In brief, ferret lung samples were processed and fluorescently labeled using standard methods (7). Samples were analyzed on the lectin microarray (92 probes, **Supplemental Table 1**) against a reference mixture consisting of all samples labeled with an orthogonal fluorophore. A heatmap of the normalized data, ordered by severity is shown in **Figure 1a.**

**Figure 1:**
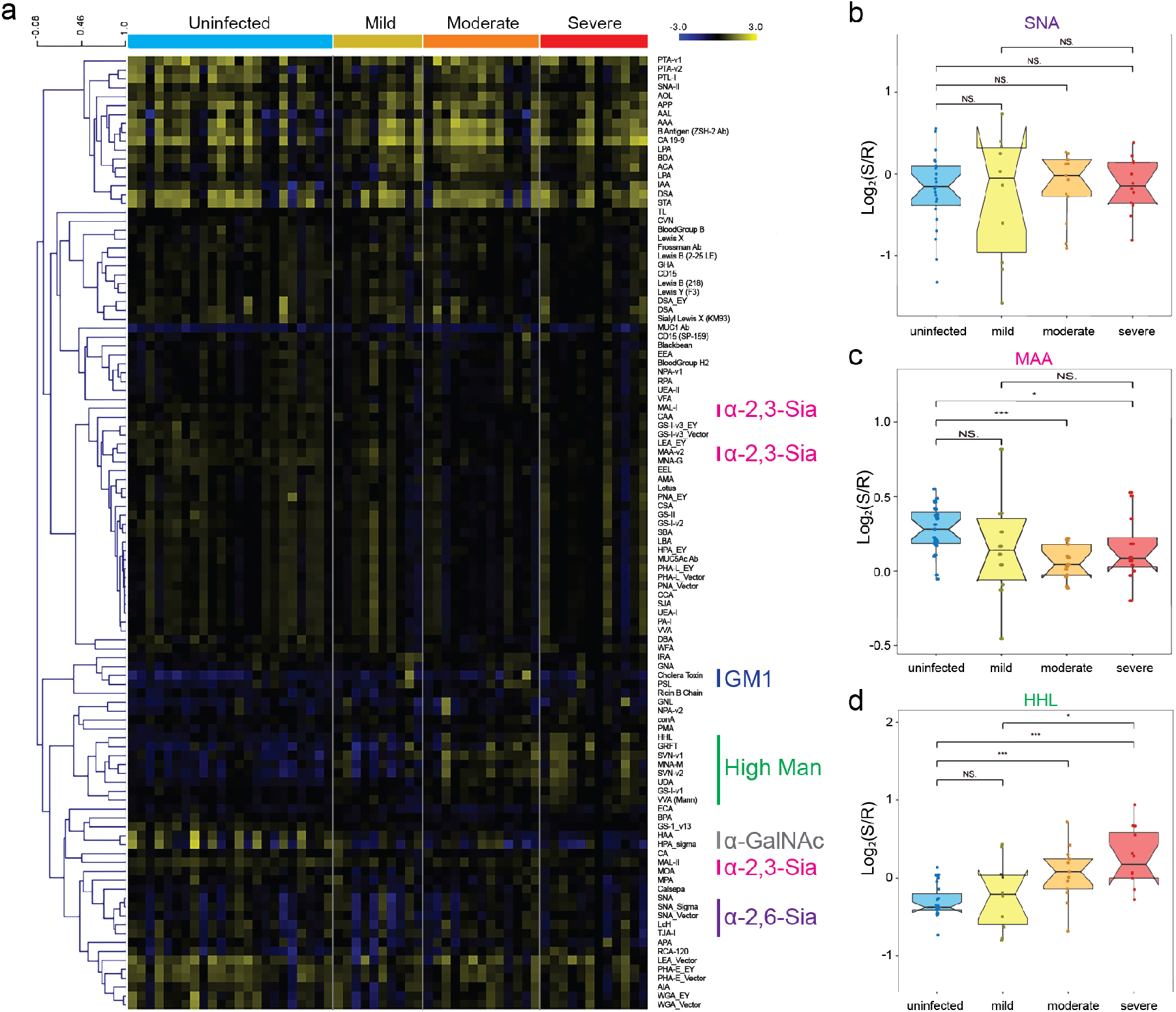
Host glycosylation change in response to influenza infection. a) Heat map of lectin microarray data. Median normalized log_2_ ratios (Sample (S)/Reference (R)) of ferret lung samples were ordered by severity (uninfected, n=4 ferrets, 4 samples per ferret; d8 infected, n=19 ferrets (mild, n=5; moderate, n=8; severe, n=6; 2 samples per ferret). Yellow, log_2_(S) > log_2_(R); blue, log_2_(R) > log_2_(S). Lectins binding α-2,6-sialosides (purple), α-2,3-sialosides (pink), GM1 (blue), high mannose (green) and N-acetylgalactosamine (grey) are indicated on the right. b) Boxplot analysis of lectin binding by SNA (α-2,6-sialosides). c) MAA (α-2,3-sialosides) and HHL (high mannose) as a function of severity. (uninfected:blue, mild: yellow, moderate: orange, high: red). N.S.: Not statistical, **: *p* < 0.01, ***: *p* < 0.001, Wilcoxon’s t-test.

Given the importance of sialic acid to influenza virus biology, we examined our data to determine whether either α-2,6-linked (lectins: SNA, TJA-I, **Fig. 1b**, **Supplemental Fig. 1a**) or α-2,3-linked sialic acid levels (MAA, MAL-I, MAL-II, **Fig. 1c**, **Supplemental Fig. 1b**) were responsive to infection. We observed no changes in α-2,6-sialic acid, the target of the H1N1pdm09 HA (21). However, we saw a subtle, yet statistically significant, decrease in α-2,3-sialic acid (~11-14% loss), which is cleaved by the viral neuraminidase (NA) (22, 23). The ganglioside GM1, a known lipid raft marker that co-localizes with HA in influenza-infected cells (24), increased in abundance upon infection as indicated by the binding of cholera toxin subunit B (~21% increase, **Supplemental Fig. 2a**). We also observed a loss of α-GalNAc (HAA, HPA, 28-34% loss, **Supplemental Fig. 2b**), an epitope predominantly detected on mucins. None of these observed changes in the host glycome correlated with severity.

However, we did observe a severity-dependent glycomic signature: lectins that bound high mannose (HHL, GRFT, SVN, UDA, **Fig. 1c, Supplemental Fig. 3**) showed increased binding in a severity dependent manner. High mannose is an early product of the *N*-glycan pathway, predominantly seen as an intracellular epitope (12). These lectins showed a statistically significant ~47-50% increase in binding to lung tissue from severely infected animals compared to uninfected controls. In contrast, ferrets with mild infections displayed a smaller increase in high mannose levels (~12-19%, dependent on lectin).

Overall, our lectin microarray data argues that although we see changes in sialic acids with infection, they have no correlation with severity. Rather, high mannose, an epitope not typically associated with influenza pathogenesis, appears to directly correlate with severity.

### Histopathology Shows High Mannose Associates with Alveolar Severity

As the influenza virus infection progresses to a more severe state, it moves from the upper respiratory tract into the alveolar cells of the lungs. The earliest responses are seen in the infected airway or lung alveolar epithelial cells. These cells become damaged, leading to lung dysfunction and pneumonia (25). To more directly observe whether high mannose levels are associated with damage in ferret lung, we next performed lectin histology on inflated lung tissue from day 8 infected ferrets. In general, we observed a correlation between damage to the lung, as defined by consolidation and inflammatory cell infiltrate, and staining with the high mannose binding lectin HHL (**Fig. 2a**). In normal lung, HHL staining was confined to the bronchiole epithelium and basement membrane. In contrast, in infected and consolidated lung tissues we observed binding to compacted alveolar spaces and bronchial plugs. We also observed binding to inflammatory cells and sloughed bronchial epithelium. We observe a strong correlation between HHL staining and the alveolar severity score, which reflects alveolar damage and inflammation caused by the infection (**Fig. 2b**). Overall, our data shows that high mannose is associated with both direct damage to the lungs, observed by histology, and overall illness levels, observed by weight loss.

**Figure 2:**
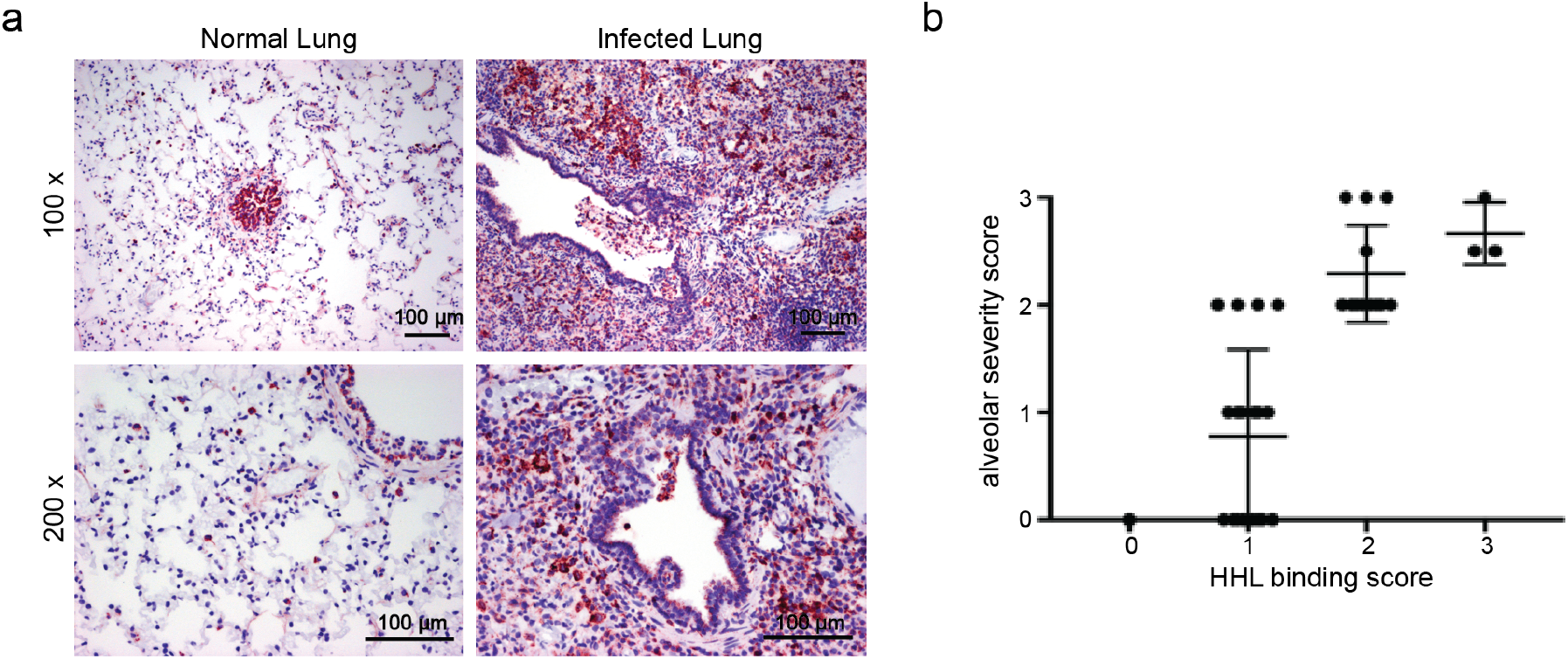
High mannose levels correlate with severity and damage. a) Lectin histology of ferret lung. Formalin-fixed paraffin embedded (FFPE) tissues stained with biotinylated HHL and avidin peroxidase (red), counterstained with hematoxylin (blue). Images shown were obtained with 10x and 20x objectives resulting in 100x and 200x magnification as labeled (factoring in 10x eyepiece lens). Scale bar: 100 μm. b) Correlation of HHL binding scores with the alveolar severity scores in histology samples. The following semiquantitative scoring system was used to evaluate HHL staining: 0, rare or occasional cells but <5% of fields; 1, >0.5 to 0.25 low-power fields; 2, >0.25 low-power fields; 3, essentially all low-power fields. Scoring of alveolar severity: 0 = normal, 1 = mild, 2 = moderate, 3 = severe. Scores for severity were averaged over fields.

### High mannose is induced early in the course of infection

Viral titers in the lungs reach high levels early in the course of influenza infection and decline significantly by day 8, with few residual viral particles by day 14 (**Supplemental Fig. 4**). At day 8 the virus is no longer replicating and is being eliminated from the lung (20). Host response to the virus is well established by this point. If glycosylation plays a role in inducing the damage leading to severity, we would expect changes to be observed early in influenza pathogenesis.

To examine whether glycomic changes in the host occur early or late in influenza pathogenesis, we performed a time-course analysis where infected ferrets were sacrificed at days 1, 3, 5, 8, and 14 (n=12 for days 1-3, n=8 for day 5, n=19 for day 8, n=8 for day 14, for a total of 59 ferrets). We analyzed 2 lung punch biopsies per ferret. The time course experiments and our previously discussed analysis were performed concurrently and analyzed on the same set of arrays. Thus, data for day 8 ferrets and control (day 0) samples are the same as in **Figure 1a**. Severity was determined by weight loss for ferrets sacrificed at days 8 and 14. Severity cannot be determined at earlier time points as the weight loss nadir has not been reached at days 1-5. Lung punch biopsies were analyzed as previously described. The heatmap for the overall analysis is shown in **Supplemental Fig. 5**.

Time course analysis revealed dynamic changes in host glycans upon infection. Levels of α-2,6-linked sialic acid increased upon infection, plateauing at day 3 (~31% change), before decreasing to baseline levels by day 8 (**Supplemental Fig. 6a**). This dynamic change in host glycans may play a role in propagating the infection as α-2,6-sialosides are the host receptor for human influenza virus, including H1N1pdm09. Levels of α-2,3-linked sialic acid declined rapidly by day 1 (~22%) and were only partially recovered at day 8 (**Supplemental Fig. 6b**). Levels of GM1 dramatically increased early in the course of infection (day 1, ~72%), perhaps due to increased viral budding from lipid rafts. These levels rapidly decreased by day 3, in line with decreased viral titers in the lung tissue (**Supplemental Figs. 5 & 6c**). Although these glycomic changes probably play a role in influenza pathogenesis, we cannot determine whether they influence severity at the early time points.

High mannose, which correlates to severity at day 8, was strongly induced in some ferrets at day 1 (**Fig. 3a**). At this early time point we are unable to predict whether ferrets with higher levels of high mannose have a more severe immunological response to influenza virus infection overall. However, HHL staining of virally infected lungs from ferrets sacrificed at day 1 correlated with damage in the lung tissue (**Fig. 3b**), similar to observations at day 8 (**Fig. 2a**). This strongly suggests that high mannose is induced immediately upon infection with influenza virus and that the levels of the epitope are directly correlated with severity even at the early time points.

**Figure 3:**
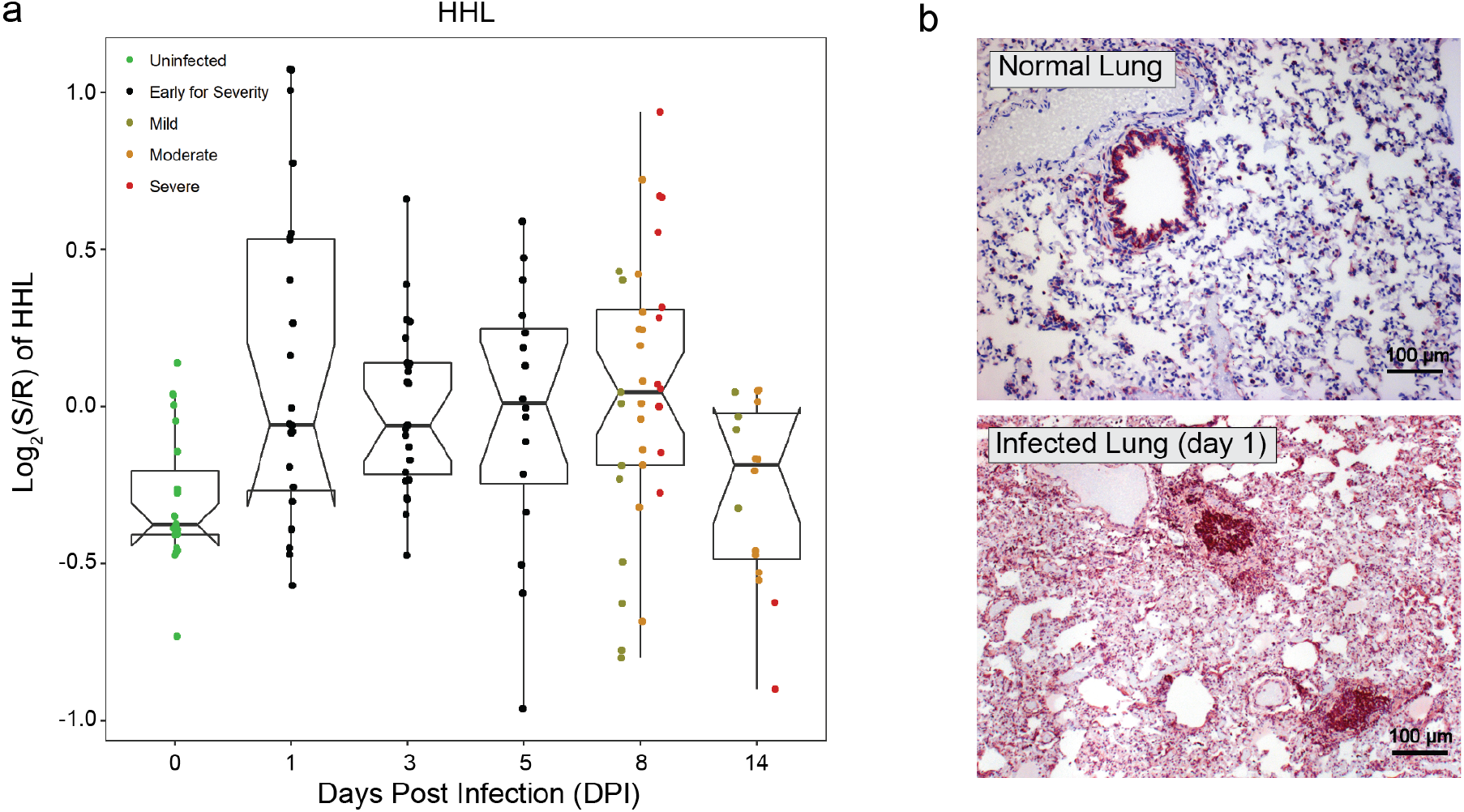
High mannose was induced at early time points in response to influenza virus infection. a) Boxplot analysis of high mannose (HHL-binding) at different time points following H1N1pdm09 virus infection (t= 0, 1, 3, 5, 8, 14 days). Median normalized log_2_ ratios (S/R) of ferret lung samples were plotted. Severity is indicated by color (green: uninfected, black: early for severity, dark yellow: mild, orange: moderate, red: severe). b) Lectin histology of ferret lung tissue. FFPE tissues were stained with biotinylated HHL developed with avidin peroxidase (red) and counterstained with hematoxylin (blue). Scale bar: 100 μm.

### Infection of Human Cell Lines with Influenza Induces High Mannose

We next tested whether the induction of high mannose upon infection was specific to our ferret model or whether it could be observed in human cell culture models as well. For our initial experiments, we used the adenocarcinoma A549 cell line, a common cell line in influenza research (26, 27), which derives from human alveolar basal epithelia. In brief, cells were infected with A/Puerto Rico/8/1934(H1N1) (PR8), an H1N1 strain commonly used in cell culture experiments. After 24 h, cells were fixed and stained for influenza nucleoprotein (anti-influenza antibody, green) and high mannose (HHL, red, **Fig. 4a)**. We observed strong induction of high mannose in infected cells. Treatment of cells with Endo H, an endoglycosidase specific for high mannose and hybrid structures (28), abolished lectin staining, as did treatment with methyl mannose (**Supplemental Fig. 7**), confirming the induction of high mannose upon infection. This epitope is induced both internally and at the cell surface, as confirmed by deconvolution microscopy (**Supplemental Fig. 8**). High mannose was observed to increase in internal structures early in the infection (8h, **Supplemental Fig. 9**) before migrating to the cell surface. To test whether the high mannose response occurs in primary human cells, we treated primary bronchial epithelia with PR8 and analyzed them as described. Again, we observed a strong increase in high mannose in response to viral infection (**Fig. 4b**). Infection of A549 cells with other influenza strains (H1N1pdm09, H3N2, influenza B, **Fig. 4c**), also resulted in an increase in high mannose, indicating that the response is not strain specific.

**Figure 4:**
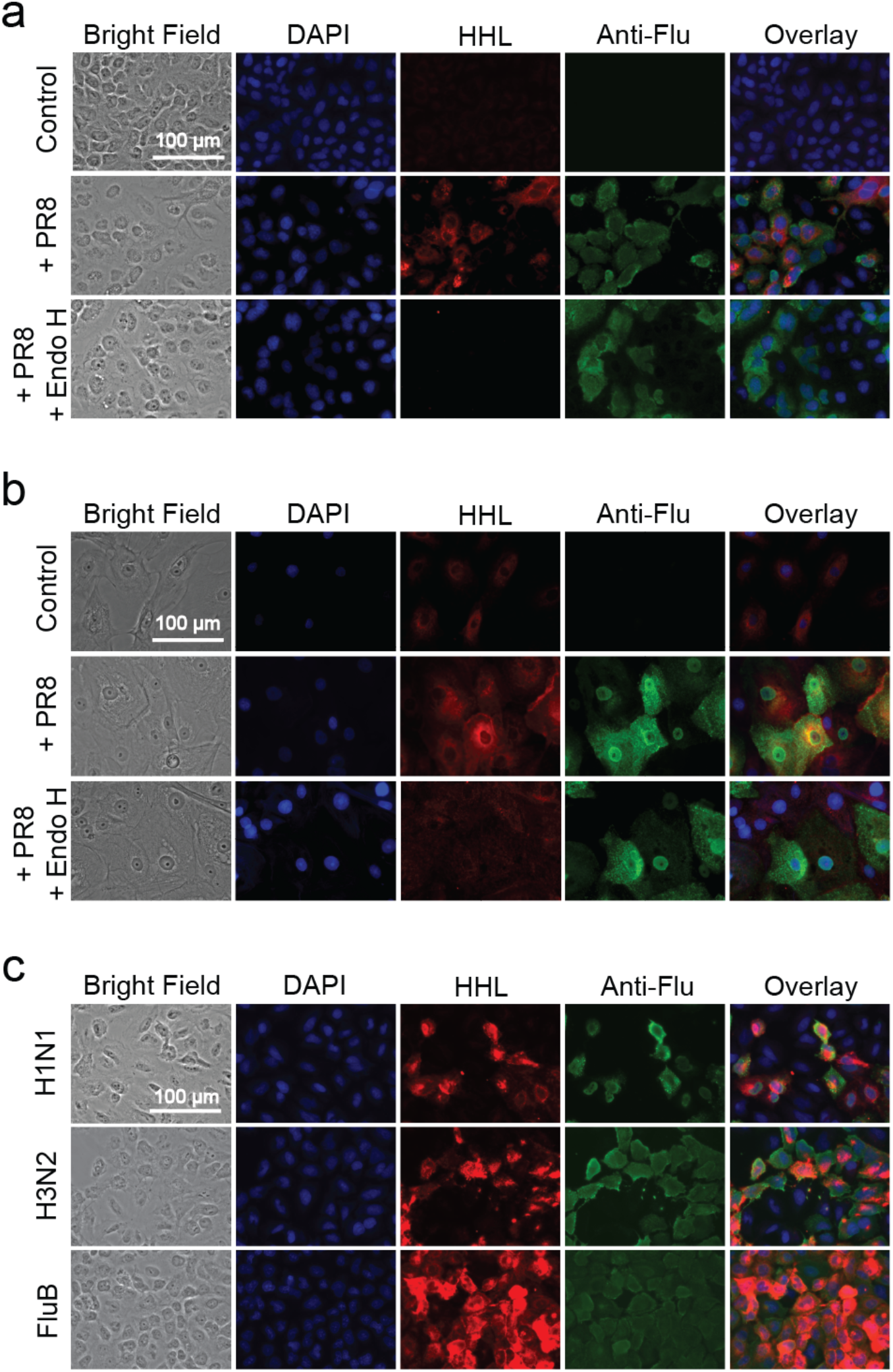
High mannose response in human A549 cells as a result of influenza virus infection. Fluorescence microscopy of a) A549 cells 24 h post infection with PR8, b) primary bronchial epithelial cells 24 h post infection with PR8 and c) A549 cells infected with additional influenza strains (H1N1pdm09, H3N2, Influenza B). All cells were co-stained with biotinylated HHL (2°: streptavidin Cy5, red), mouse anti-influenza nucleoprotein (anti-Flu) antibody (2°: anti-mouse IgG-Cy3, green), and DAPI (blue). For Endo H controls, enzyme treatment was performed prior to staining. Bright field and overlay images of DAPI, HHL and anti-Flu stained images are shown. Scale bar: 100 μm. For each experiment, three biological replicates were performed and a minimum of six images was captured. Representative images are shown.

Given that high mannose is a precursor in the N-glycan pathway, a question that arises is whether high mannose could act as an independent signal without impacting complex N-glycan levels. Recent studies have shown that the trimming mannosidases, MAN1A1 and MAN1A2 can be independently deleted without impacting complex N-glycans (29, 30). In addition, both enzymes are predicted to be highly regulated by miRNA (31), and miRNA downregulating MAN1A2 increased high mannose without decreasing core fucose, a marker of complex N-glycans (11). To test this more directly, we performed lectin microarray analysis of A549 cells infected with PR8 (**Fig. 5a, Supplemental Fig. 10**). In line with our observation in ferret lungs and our microscopy data, we again observed an increase in high mannose (GRFT, SVN, HHL, UDA). However, we did not observe a corresponding decrease in complex N-glycan epitopes (indicated by red *: branching: PHA-L, polyLacNAc: DSA, WGA, core fucose: LcH). A decrease was observed in α-2,6-linked sialic acid (SNA, TJA-II) in line with previous data on influenza infections in cell culture (32). Overall our data supports the ability of high mannose to act as a signal of infection and/or damage.

**Figure 5:**
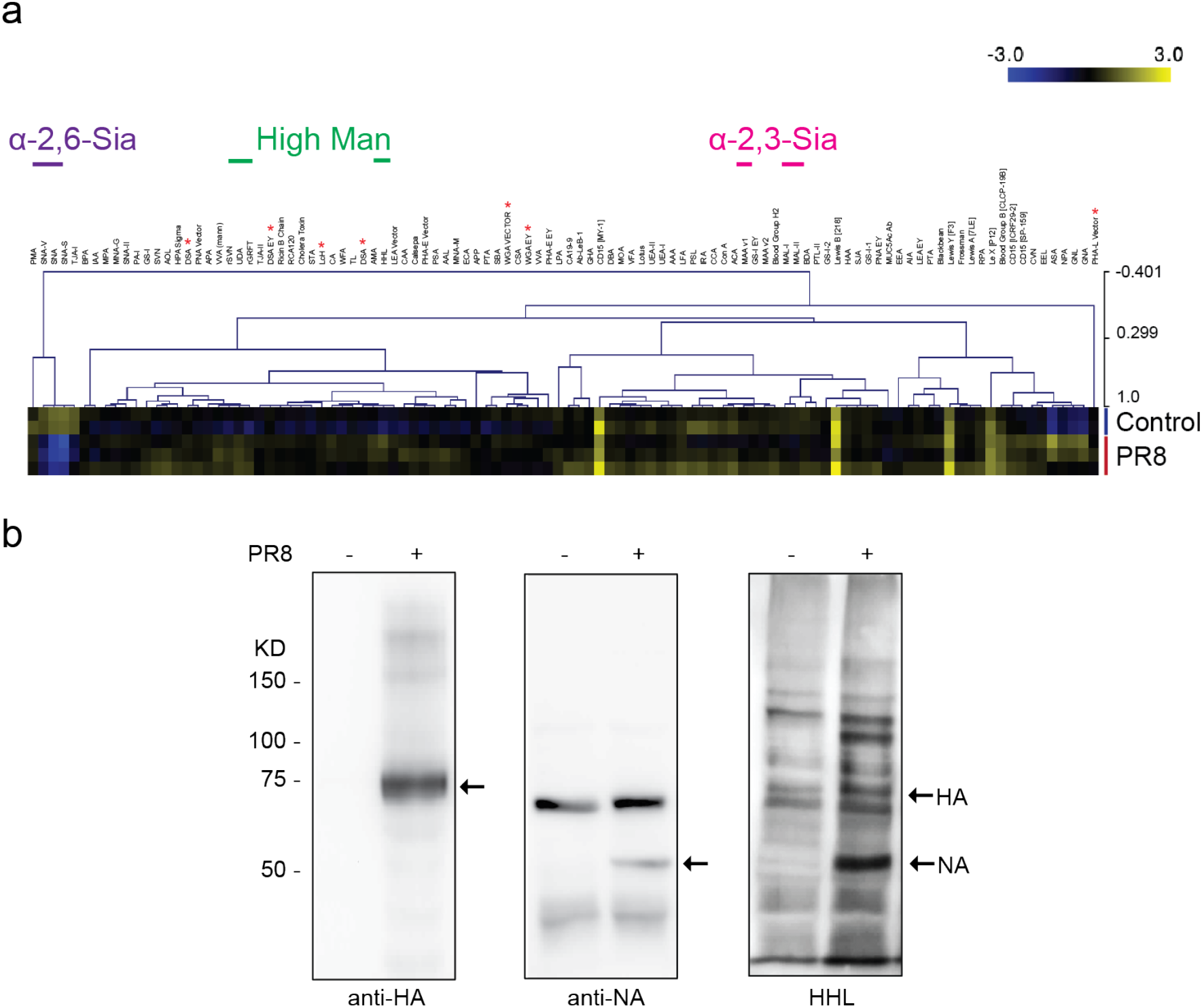
Lectin microarray and lectin blot analysis of influenza infected A549 cells. a)) Heat map of lectin microarray data. Median normalized log_2_ ratios (Sample (S)/Reference (R)) of samples from A549 cells infected with PR8 (n=3) or uninfected controls (n=2). Yellow, log_2_(S) > log_2_(R); blue, log_2_(R) > log_2_(S). Lectins binding α-2,6-sialosides (purple), α-2,3-sialosides (pink) and high mannose (green) are indicated at top. Red * indicates lectins binding complex N-glycans. Graphical representation of select lectin data is show in **Supplemental Fig. 10**. b) Western blot and lectin blot analysis of the A549 cells. Duplicate lanes were run simultaneously and transferred to nitrocellulose. Blot was then divided and stained for either influenza proteins (rabbit polyclonal anti-HA (1:1000) or rabbit polyclonal anti-NA (1:1000) and developed with anti-rabbit-HRP (1:5000)) or for high mannose (biotinylated HHL (20 μg/mL) followed by streptavidin HRP (1:5000)). HA and NA proteins are indicated by arrows. Even loading was checked by Ponceau (**Supplemental Fig. 11)**.

### Influenza Virus Induces High Mannose on Endogenous Glycoproteins

One possible source of high mannose is the influenza virus itself, which has three potential glycoproteins: HA, NA, and Matrix-2 protein (M2). HA is the major glycoprotein of influenza. *N*-glycosylation of HA is important in receptor binding, immune response, and viral stability (33). Glycosylation of NA is essential for the enzyme activity (33). M2, a small transmembrane protein of about 11 KD, is at 10-100 fold lower abundance than HA on the virus surface and has a putative *N*-glycosylation site at which glycosylation is not commonly observed (33–36). To determine whether the increase in high mannose was due to influenza virus glycoproteins or to changes in glycosylation of host proteins, we performed lectin blot analysis in tandem with Western blot analysis of HA and NA, the major influenza virus glycoproteins (**Fig. 5b, Supplemental Fig. 11**). Upon infection, we detect increased levels of high mannose across multiple proteins, both influenza and host, as determined by HHL binding. We observed a strong increase in high mannose for a band correlating to NA (indicated by arrows, **Fig. 5b**). Quantitation of the lectin blot, excluding presumed HA and NA proteins, reveals a 120% increase in HHL binding, indicating a major increase in high mannose on host proteins. Our data shows that high mannose induced in human cells is an endogenous host response to influenza virus infection.

### High Mannose is Induced via the IRE1/XBP1 Pathway

Recent work provides evidence for the involvement of the unfolded protein response (UPR) in determining influenza severity. Influenza virus activates the IRE1 arm of the UPR pathway, inducing the active form of the transcription factor XBP1 (XBP1s) (37). Influenza severity and lung injury in mouse models directly correlate with the degree of induction of this pathway (13). Recent studies have shown that activation of XBP1s in cells can alter glycosylation in a cell type dependent manner, providing a potential link between influenza infection and the observed changes in host glycosylation (38, 39). We first examined whether influenza infection activated the UPR pathway in our ferret model. Transcriptomic analysis revealed upregulation of key UPR markers HSP90B1 (also known as Grp94) and HYOU1 (40) in ferret lung on day 1 post-infection (**Fig. 6a**). This indicates that UPR is rapidly induced by influenza in ferrets, in line with previous observations in other systems (13, 37). To test whether there is a direct link between UPR activation by influenza virus and the induction of high mannose, we used the established UPR inhibitor 4μ8C (4-methyl umbelliferone 8-carbaldehyde) in our cell culture model. This inhibitor prevents splicing of XBP1, the key step in activation of XBP1s by IRE1 (41). Inhibition of UPR in PR8-infected A549 cells by 4μ8C (64 μM) prevented high mannose induction by the virus (**Fig. 6b**, **Supplemental Fig. 12a**). This indicates that induction of the high mannose epitope occurs downstream of the XBP1s arm of the UPR. Lectin microarray analysis revealed 4μ8C inhibited high mannose but did not impact the change in sialylation observed upon infection, arguing that the impact of inhibiting this pathway is specific to the high mannose response (**Supplemental Fig. 12b**).

**Figure 6:**
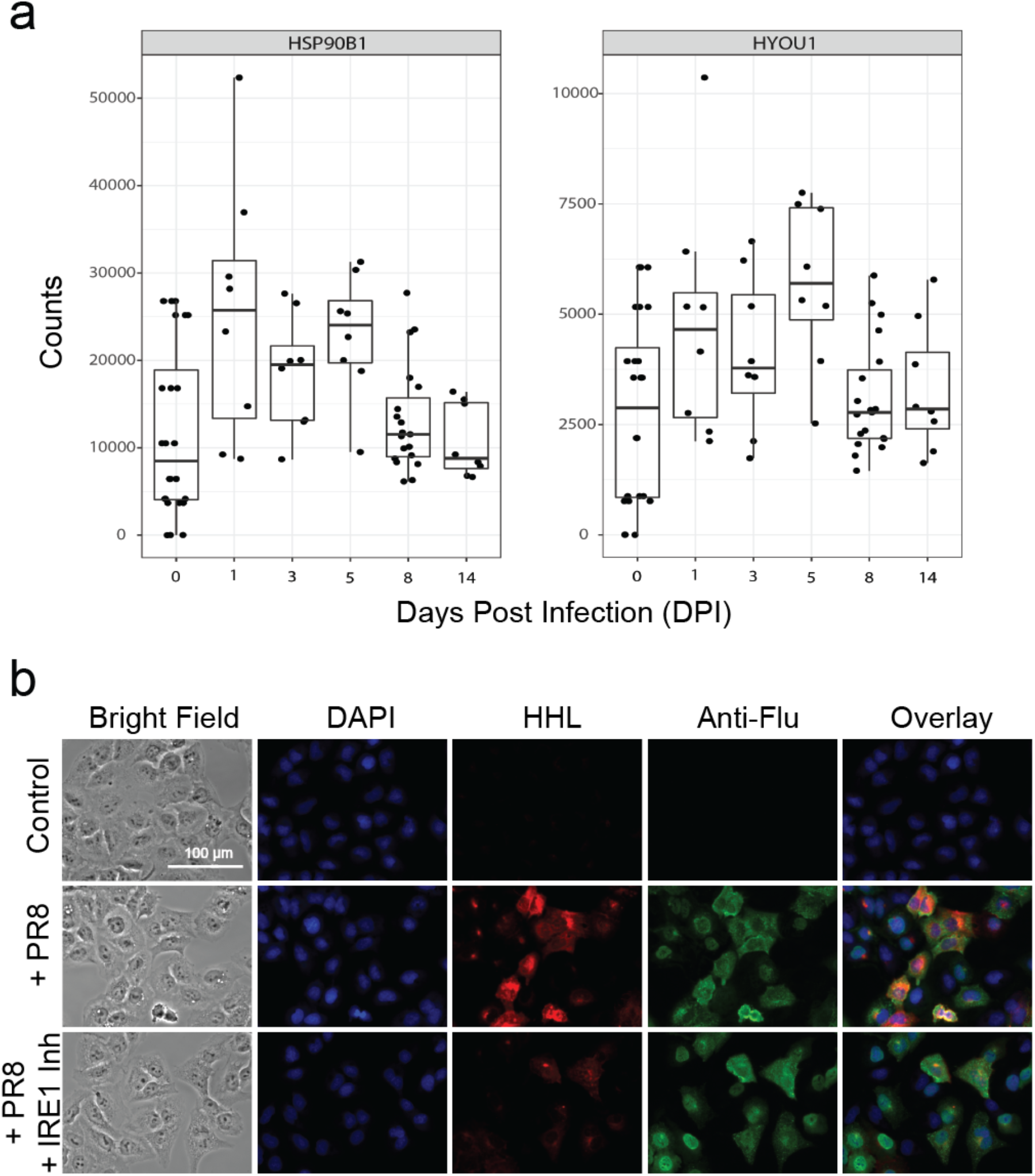
High mannose is induced via the unfolded protein response. a) Boxplot analysis of transcript levels of UPR markers HSP90B1 and HYOU1 in lung tissues from H1N1pdm09 infected ferrets from time course analysis. b) Fluorescence microscopy of A549 cells treated with IRE1 inhibitor 4μ8C (24 h) prior to infection. Cells were fixed, stained and imaged as previously described. Scale bar: 100 μm. For each experiment, three biological replicates were performed and a minimum of six images was captured. Representative images are shown. Lectin microarray analysis of A549 cells treated as in b is shown in **Supplemental Figure 12b**.

### Innate Immune Activator MBL2 Recognizes Infected Cells via High Mannose

Currently, there is an emerging consensus that over-activation of the innate immune response may be a dominant cause of lung injury and influenza severity (20, 25). Carbohydrate recognition plays a significant role in innate immunity mediated by a series of innate immune lectins including the dendritic cell-specific intercellular adhesion molecule-3-grabbing non-integrin (DC-SIGN), human DC immunoreceptor (DCIR), langerin, dectin-2, mincle, mannan-binding lectin (MBL2), and surfactant proteins A and D (SP-A/D) (12, 42). Innate immune lectins are thought to protect against infection through recognition of foreign pathogens. We focused our attention on MBL2. MBL2 binds influenza viral particles and was originally ascribed a protective role (43). However, knockout of MBL2 in mouse mitigated the severity of influenza infection, rather than increasing viral load as originally theorized (18). Upon binding, MBL2 activates the complement cascade and corresponding inflammatory response (44). A recent study on critically ill patients infected with H1N1pdm09 virus showed a strong correlation between levels of MBL2 in the blood and patient death (19). In addition, schizophrenics and multiple sclerosis patients, both have high levels of MBL2 and unusually high death rates from influenza and pneumonia (45–48). These studies strongly suggest that MBL2 plays a determining role in the severity of response, but how this occurs is unknown.

MBL2 recognizes several glycan epitopes including Lewis structures (14, 15), high mannose (16), yeast mannans (17) and fucose (15). Given that high mannose is a ligand for MBL2, we wanted to test whether MBL2 would recognize influenza virus-infected human cells. We incubated infected and control A549 cells with recombinant human MBL2 and examined binding by fluorescence microscopy (**Fig. 7**). Uninfected A549 cells were not recognized by MBL2. In contrast, infected cells were strongly recognized by the innate immune lectin. This data is in keeping with previous work that demonstrated complement-dependent lysis of influenza-infected hamster cells mediated through binding of guinea pig mannan-binding lectin (49).

**Figure 7:**
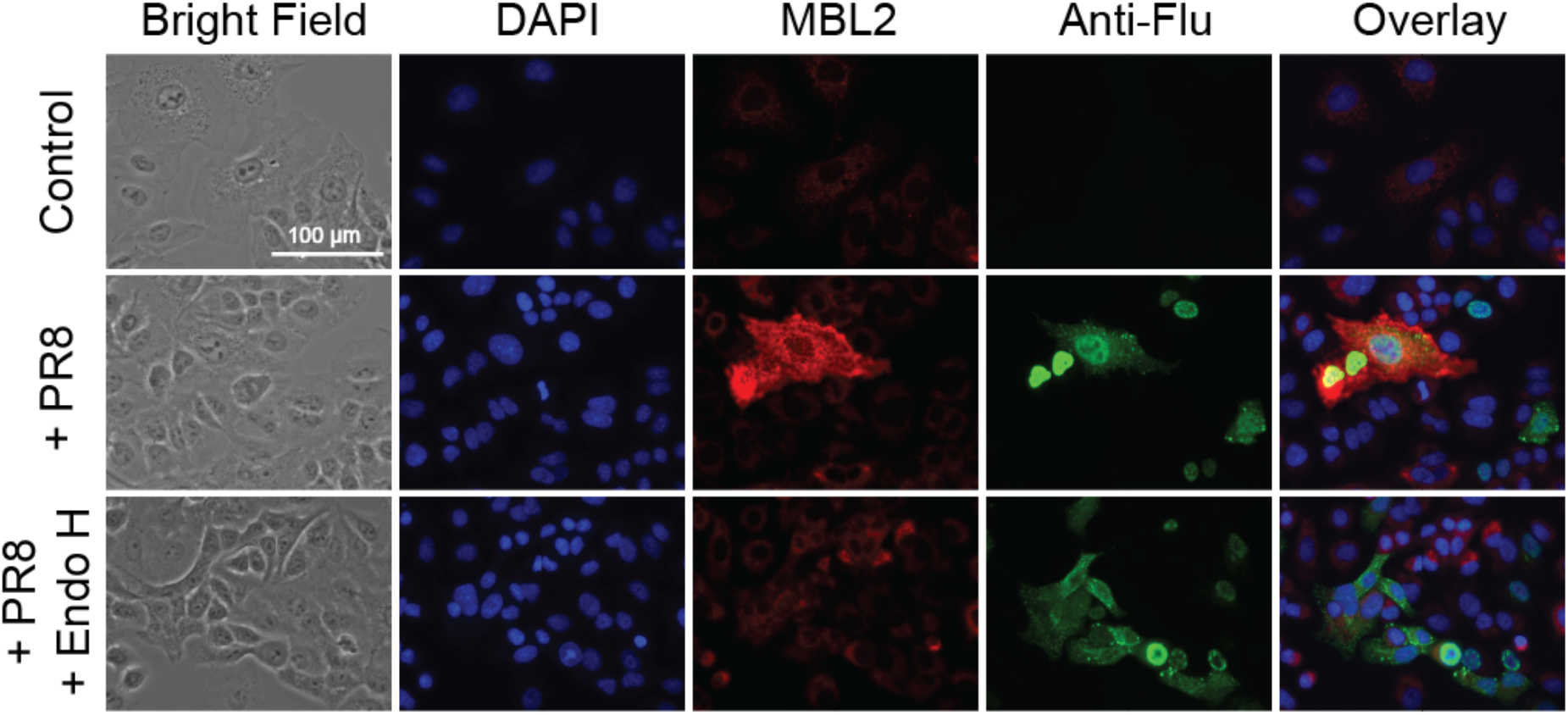
Mannose is recognized by mannose binding lectin (MBL2). MBL2 staining of A549 cells. A549 cells (PR8 infected (24 h), and Control) were incubated with recombinant human MBL2 and stained with mouse anti-MBL (Biotin) antibody (2°:Cy3 Streptavidin, red). Cells were counterstained for infection (1°: mouse anti-Flu labeled with Cy3, green), and DAPI (blue). Scale bar: 100 μm. For each experiment, two biological replicates were performed and a minimum of six images was captured. Representative images are shown.

To test whether recognition is dependent upon the high mannose epitope, we removed all high mannose and hybrid N-glycans using Endo H. Removal of these glycans abolished the signal, providing evidence that the recognition is via the high mannose epitope (**Fig. 7**). Our data suggest a potential role for high mannose in recruiting MBL2 to host tissue in influenza virus infection; this could contribute to severity through activation of the complement cascade and corresponding inflammatory response.

### Conclusions

Lectin-glycan interactions play a major role in infectious disease, impacting all stages from colonization to disease progression and host response (50–52). Current studies of these interactions almost exclusively focus on their roles in invasion or on innate immune lectin recognition of pathogen glycans (50). In contrast, little is known about change in the host glycome upon pathogen invasion and how this might impact disease progression.

Herein, we use a ferret model of influenza virus infection in tandem with our lectin microarray technology (6, 8) to study the host response to the 2009 H1N1 influenza virus and its relationship to disease progression. Our data suggest that induction of high mannose upon influenza virus infection is a key mediator of severity. Based on our work and the current literature, we hypothesize that activation of the unfolded protein response (UPR) upon influenza infection turns on expression of high mannose, which is then recognized by the innate immune lectin MBL2, activating the complement cascade and subsequent inflammation (**Fig. 8**). Inhibition of the complement cascade with C3a inhibitors reduces damage in H5N1 influenza virus-infected mice, providing additional evidence for our hypothesis (53). Our results indicate that high mannose can be induced through the UPR upon influenza infection (**Fig. 6, Supplemental Fig. 12**). The use of high mannose by the cell, as a type of damage or distress signal, makes sense as it is not detected at the cell surface on most cell types, but is recognized by a significant number of innate immune lectins, including MBL2 (**Fig. 7**) (12). In addition, high mannose has the potential for independent regulation from the complex N-glycan pool (**Fig. 5a**) (11, 29–31). Taken together, our work provides a new mechanism that could explain the observation that both UPR activation and MBL2 levels directly associate with severity of outcome from influenza infection (13, 18, 19).

**Figure 8:**
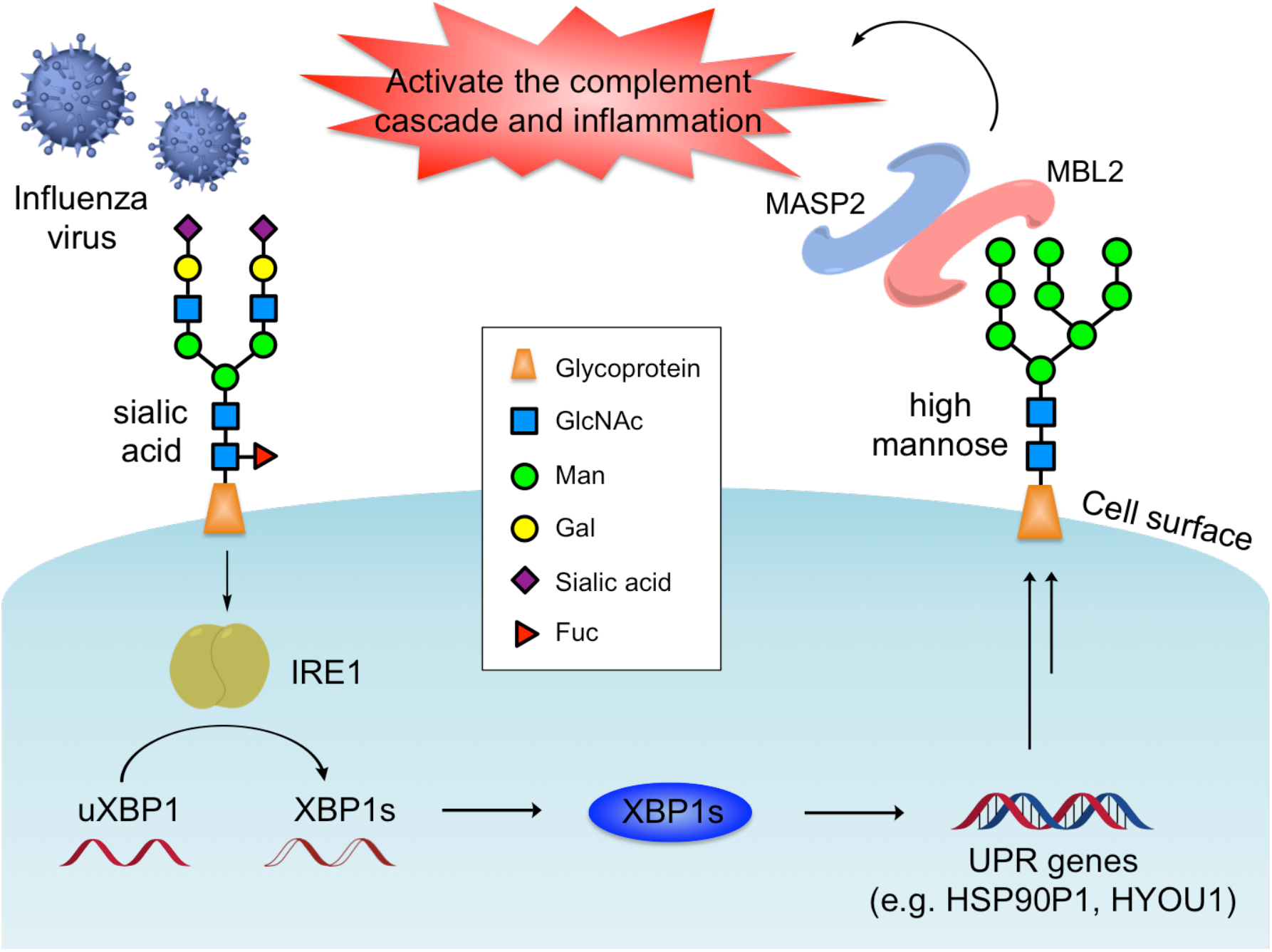
Schematic representation of proposed mechanism. Influenza infection activates the IRE1 arm of the UPR, leading to the production of high mannose on the cell surface. High mannose is recognized by the innate immune lectin MBL2, which in turn activates the complement cascade and accompanying immune response, determining severity.

A picture is emerging in which high mannose could act as a signal to our immune system through MBL2 (and potentially other mannose-binding lectins) to clear damaged and infected cells. This hypothesis, which requires further testing, could be relevant for not only influenza but also other respiratory viruses where the innate immune system plays a role in severity, such as SARS-CoV-2 (54). It predicts that an overabundance of high mannose (as seen in the ferrets) or high MBL2 (as seen in patients) can dysregulate the immune system, leading to severe damage and death. Notably, two populations that have high levels of activated MBL2, schizophrenics and multiple sclerosis patients, have unusually high death rates from influenza and pneumonia (45–48). Our work provides a potential new pathway for intervention in influenza virus infection that could spur the development of therapies that would make influenza virus-induced disease no more deadly than the common cold.

## METHODS

### Influenza virus and infection of ferrets

Fitch ferrets (*Mustela putorius furo*, female) were obtained from Triple F Farms (Sayre) and verified as negative for antibodies to circulating influenza A (H1N1 and H3N2) and B viruses. Adult ferrets were defined as 6-12 months of age. Ferrets were pair housed in stainless steel cages (Shor-Line) containing Sani-Chips laboratory animal bedding (P.J. Murphy Forest Products) and provided with food and fresh water ad libitum. Ferrets were administered intranasally the H1N1pdm09 virus, A/California/07/2009, at a dose of 106 PFU. The animals were monitored daily for weight loss and disease symptoms including elevated temperature, low activity level, sneezing, and nasal discharge. Animals were randomly assigned to be sacrificed at day 1, 3, 5, 8, or 14 post-infection (DPI) or euthanized if their clinical condition (e.g., loss of >20% body weight) required it. Blood was collected from anesthetized ferrets via the subclavian vena cava post-infection. Serum was harvested and frozen at −20 ± 5°C. After serum was collected, necropsies were performed to collect lung tissue. Severity of infection was determined for all ferrets studied who were sacrificed at day 8 or later (n=45) using quartiles to define populations. The lowest quartile was defined as mild (weight loss less than 10.5%), the middle quartile as moderate (weight loss between 10.5% and 16.2%), and the highest quartile as severe (weight loss greater than 16.2%).

### Lectin microarray

Ferret lung tissue samples were washed with PBS supplemented with protease inhibitor cocktails (PIC) and sonicated on ice in PBS with PIC until it was completely homogenous. Then the homogenized samples were prepared and Alexa Fluor 647-labeled as previously described (7). Reference was prepared by mixing equal amounts (by total protein) of all samples and labeled with Alexa Fluor 555 (Thermo Fisher). **Supplemental Table 1** summarizes the print list and buffers. Printing, hybridization, and data analysis were performed as previously described (7).

### Lectin histochemistry

Formalin-fixed paraffin embedded (FFPE) sections (5 μm) containing left upper and lower lung lobes were cleared in Histoclear (3 × 5 min) (National Diagnostics). Sections were rehydrated with graded alcohols as follows: 100% ethanol (2 × 5 min), 95% ethanol (1 × 5 min), 70% ethanol (1 × 5 min), and H_2_0 (1 × 5 min). To inactivate endogenous peroxidases, rehydrated sections were immersed in 3% H_2_O_2_/70% methanol solution (Fisher Scientific) for 30 min. Sections were incubated with biotin-conjugated HHL (1:300, Vector Laboratories) overnight at 4°C. After washing in PBS, sections were incubated with an avidin/biotin-based peroxidase system (VECTASTAIN Elite ABC HRP Kit, Vector Laboratories) followed by substrate deposition with VECTOR NovaRED Peroxidase (HRP) Substrate Kit (Vector Laboratories). Nuclear counterstain was performed with Gill’s Hematoxylin (American MasterTech Scientific) followed by mounting with Permount (Fisher Scientific).

### Cell culture and infection assays

The A549 cell line (ATCC) was grown in media [F-12K medium with 10% (vol/vol) FBS (Atlanta Biologicals)] at 37°C in 5% CO_2_. For the infection assay, A549 cells were seeded at a density of 250,000 cells in a 35 mm glass-bottom dish and allowed to grow under standard culture conditions (37°C, 5% CO_2_); 24 h later the dish was washed with PBS and treated with PR8 virus (MOI =2) in the infection medium [F-12K with 2% BSA fraction V (Thermo Fisher), 1% antibiotic-antimycotic (Thermo Fisher), and 1 μg/ml TPCK-treated trypsin (Worthington Biochemical)] for 1 h (37 °C, 5% CO_2_). The cells were washed with the infection medium and incubated in the infection medium for 8, 6, or 24 h, as indicated prior to harvesting. Control cells were treated identically, however no PR8 virus was added to the media. <u>IRE1 inhibition</u>: Cells were seeded at a density of 120,000 cells in 35 mm glass-bottom dishes and grown in standard condition. After 24 h cells were treated with either IRE1 inhibitor 4μ8C (64 μM) or DMSO (Control). After an additional 24 h, cells were treated with PR8 virus as described above. For IRE1 inhibitor treated cells, 4μ8C (64 μM) was maintained in the culture media throughout the infection. All cells were then either lysed for Western blot analysis, processed for lectin microarray analysis (7), or fixed and stained for fluorescence imaging.

### Fluorescence microscopy

Cells were fixed with 4% paraformaldehyde in PBS (30 min, RT), washed 3 × with PBS, and permeabilized with 0.2% Triton-X in PBS (5 min). Fixed cells were washed with PBS and blocked for 1 h (PBS, 1% BSA) at 37°C (5% CO_2_). Cells were stained with both biotinylated HHL (20 μg/ml in 10 mM HEPES buffer; Vector Laboratories) and mouse anti-influenza A (1 μg/ml in 10 mM HEPES buffer; Abcam) for 1 h at 37°C. After washing 3 × with PBS, cells were stained with Cy5-streptavidin (10 μg/ml; ThermoFisher) and anti-mouse IgG Cy3 (5 μg/ml; 1 h, 37 °C; Abcam) in PBS for 1 h (37°C, 5% CO_2_). After washing 3x with PBS, the cells were stained with DAPI (600 nM; 5 min, RT; ThermoFisher) prior to imaging. Samples were imaged using a 40× PlanFluor objective, NA 0.3 and a Quad filter cube (DAPI, FITC, Texas Red, Cy5) on an Eclipse TE 2000-U microscope (Nikon). A minimum of six images were obtained for each sample in each channel. Samples stained with the same lectin or antibodies were imaged under identical conditions. <u>Endo H controls</u>: Cells were treated with Endo H (NEB, 1:10 in glycobuffer 3; 1h, 37 °C, 5% CO_2_) prior to HHL staining. <u>Mannose inhibition</u>: biotinylated HHL was incubated with methyl mannose (200 mM; 30 min, RT; Sigma) prior to HHL staining.

### MBL2 staining of A549 cells

A549 cells were cultured and fixed and blocked as previously described. Cells were stained with MBL2 [10 μg/ml in MBL2 binding buffer (20 mM Tris, 1 M NaCl, 10 mM CaCl_2_, 15 mM NaN_3_, 0.05% triton X-100, pH 7.4); 1h, 37 °C; Abcam]. After washing with MBL2 wash buffer (10 mM Tris, 145 mM NaCl, 5 mM CaCl_2_, 0.05% tween-20, pH 7.4) 3 ×, cells were stained with mouse monoclonal anti-MBL (Biotin) (2 μg/ml in MBL2 wash buffer; 1 h, 37 °C; Abcam) and mouse monoclonal anti-influenza A that had been labeled with Cy3 dye and dialyzed following the manufacturers protocol (1 μg/ml in MBL2 wash buffer; 1 h, 37 °C; Abcam). Cells were washed 3 × (MBL2 wash buffer) and stained with both Cy5 Streptavidin (10 μg/ml; Vector Laboratories) in MBL2 wash buffer for 1 h at 37 °C (5% CO_2_). Cells were then washed 3 × (MBL2 wash buffer) and stained with DAPI (600 nM in MBL2 wash buffer; 5 min, RT; ThermoFisher) before imaging. Samples were imaged by fluorescence microscopy as described above. Endo H treated controls were prepared as described above.

## Acknowledgements

The University of Georgia Institutional Animal Care and Use Committee approved all experiments under the Animal Use Protocol #2015-04-007, which were conducted in accordance with the National Research Council’s Guide for the Care and Use of Laboratory Animals, The Animal Welfare Act, and the CDC/NIH’s Biosafety in Microbiological and Biomedical Laboratories guide. We thank Dr. Matthew D. Shoulders at M.I.T. for advice and Boval Biosolutions for lyophilized protease- and IgG-free bovine serum albumin (no. LY-0081). We thank Dr. Peter Palese and Dr. Adolfo Garcia-Sastre for providing the PR8 RGs plasmids. This work was supported by NIAID/NIH U01 AI111598 to E. Ghedin, B. Zhang, and L. Mahal. Publication of this research was supported, in part, thanks to funding from the Canada Excellence Research Chairs Program (L. Mahal).

