## Supplemental Information for "Glycomic analysis of host-response reveals high mannose as a key mediator of influenza severity"

***Supplemental Methods.***

**Reagents.** We purchased A549 cells, primary bronchial epithelial cells, and F12K media from ATCC, the reverse transcription kit from AB Applied Biosystems, polymerases from NEB and chemicals from Sigma-Aldrich. Lectins were purchased from E. Y. Laboratories or Vector Laboratories with the following exceptions: recombinant cyanovirin, SVN and griffithsin were gifts from B. O'Keefe (Frederick National Laboratory for Cancer Research, Frederick, MD); recombinant Gaf-D, PA-IL, PA-IIL, and RS-IIL were made as previously described (1); TJA-I and TJA-II were from NorthStar Bioproducts. Antibodies were purchased from Abcam (anti-MBL: ab203303; anti-influenza A: ab20343). Human recombinant MBL2 protein was purchased from Abcam (ab151947). The H3N2 strain used was A/New York/238/2005. The FluB strain used was B/Brisbane/60-10/2010.

**Ferret Tissue preparation.** Lungs were rinsed with cold PBS via catheterized trachea to collect bronchioalveolar lavage fluid (BAL). The right upper and lower lobes were removed and each lobe was sectioned into quadrants. Sections were snap frozen. The left upper and lower lung lobes were formalin perfused. After fixation, tissue was paraffin-embedded and 5 mm-thick sections were prepared for histopathological analysis.

**Histopathological analysis.** All lungs were inflated for this study. Tissue sections containing lobe from the left upper lung, and lobe from the left lower lung were stained with hematoxylin and eosin. Sections were scored by two blinded readers for bronchial and alveolar severity. Regional severity of bronchial (bronchi and bronchioles) and alveolar spaces based on inflammation, erosion and alveolar immune infiltration was assessed by the following scoring guidelines: 0 = normal; 1 = mild; 2 = moderate; 3 = severe (2). To ensure accurate scores for these tissues, we first flagged the slides that were not well inflated to account for potential artifacts due to poor inflation when scoring. After quantitation, we performed additional stains with anti-myeloperoxidase (MPO) to determine extent of inflammation. Normal lungs that were not well inflated have limited MPO staining, and inflamed lungs show considerable MPO staining. We then compared these scores with both our H&E scores and our HHL scores to confirm the validity of the original scores.

**Viral plaque assay.** Plaque assays were performed to determine viral burden in lung tissue. Lung tissue supernatants were obtained from frozen lung pieces that were gently thawed on ice, forced through a cell strainer (70 mm) and syringe plunger in PBS, then spun down (2500 rpm, 5 min, 4°C) to collect supernatant. Nasal washes and lung supernatants were diluted in Iscove's modified Dulbecco minimum essential medium (iDMEM). Madin-Darby canine kidney (MDCK) cells were plated ($5x10^5$) in each well of a six-well plate. Samples were diluted (final dilution factors of 100 to $10^{-6}$) and overlaid onto the cells in 100 µl of Dulbecco modified Eagle medium (DMEM) supplemented with penicillin-streptomycin and incubated for 1 h. Samples were removed, cells were washed twice, and medium was replaced with 2 ml of L15 medium plus 0.8% agarose (Cambrex, East Rutherford, NJ, USA) and incubated for 72h at 37°C with 5% CO2. Agarose was removed and discarded. The cells were fixed with 10% buffered formalin and then stained with 1% crystal violet for 15 min. Following thorough washing in distilled water (dH2O) to remove excess crystal violet, the plates were dried, number of plaques was counted, and the number of PFU per milliliter was calculated.

**Western blot and lectin blot analysis.** Cells were washed with 1x PBS twice and lysed in cold RIPA buffer (ThermoFisher, 89900) supplemented with 1000x DMSO and aqueous protease inhibitor cocktails diluted to 1x. Samples were sonicated (80% amplitude; 10 s on 10 s off for four times on ice) and centrifuged (14,000 xg, 4 °C, 30 min) to remove the debris. Equal amounts of proteins (50 µg) were resolved by 7% SDS-PAGE gel with 4% stacking, transferred onto nitrocellulose membranes, and blocked in blocking buffer (5% BSA in PBST (PBS, pH 7.4, 0.05% Tween-20), 1 h, RT). For western blot, antibodies were diluted in blocking buffer. Primary antibodies: rabbit polyclonal anti-HA (1:1000; ThermoFisher, PA5-34929), rabbit polyclonal anti-NA (1:1000; Abcam, ab21304); 4 °C overnight. The blot was washed with PBST three times then stained with secondary antibodies (anti-rabbit HRP; 1 h, RT; Abcam) in blocking buffer. For lectin blots, biotinylated HHL (20 µg/ml; 1 h, RT; Vector Laboratories) was diluted in HEPES buffer (10 mM HEPES, 115 mM NaCl, 1.2 mM $CaCl_2$, 1.2 mM $MgCl_2$, 2.4 mM $K_2HPO_4$, pH 7.4). The blot was washed with PBST three times followed by staining with streptavidin HRP in HEPES buffer described above (1:5000, 1 h, RT; ThermoFisher). Blots were developed using SuperSignal West Pico (ThermoFisher).

**mRNA-seq library preparation and data processing.** RNA was extracted from ferret whole blood with the Mouse RiboPure-Blood RNA Isolation Kit (Ambion) according to the manufacturer's protocol and incorporating a DNase treatment (Qiagen) after passing the sample through the filter cartridge. Following elution, all RNA was stored in LoBind 1.5mL tubes (Eppendorf) and a 5 µL aliquot was made in order to measure sample yield and integrity. Representative samples from each set of extracts were quantified with the Qubit BR RNA Assay (ThermoFisher Scientific) on a Qubit 2.0 Fluorometer (ThermoFisher Scientific) and their fragment size distribution was measured with an RNA Screentape (Agilent) on a TapeStation 2200 (Agilent). Sample yield ranged from 1.2 µg to 6.15 µg, but all samples had sufficient input mass for downstream library prep and RIN values were consistently >8.5. In order to meet the minimum input concentration requirements for library prep, all samples were concentrated using the ZR-96 RNA Clean and Concentrator (Zymo) in the predetermined 96-well plate format. Two processing controls (i.e. with elution buffer as input) were included during this step, one for each plate of the ZR-96 kit. Following elution, the concentration of all samples was measured using the Quant-IT RNA Assay Kit (ThermoFisher Scientific) on an M200 Microplate Reader (Tecan). Libraries were prepared using the TruSeq RNA Sample Preparation Kit v2 (Illumina) according to the manufacturer's protocol using either 1 µg of input RNA or, in the case of the controls and several low-yield samples, the maximum possible input mass. Libraries were barcoded with the supplied index adapters in a rotating scheme, which ensured that each pool had a unique set of barcodes used in combination. Each library was quantified by qPCR with the Library Quantification Kit for Illumina (KAPA Biosystems) on a Roche 480 Lightcycler (Roche) and pooled with equimolar input at 20 µM into sets of 12 or 15 libraries for sequencing on a HiSeq2500 in Rapid Run or High Output mode, respectively. For those control libraries that did not exceed 20 µM, the maximum possible input molarity was used. Each pool was re-quantified by qPCR and its fragment size distribution was measured with a D1000 Screentape (Agilent) on a TapeStation 2200 before dilution to 2 nM. Each pool was further diluted to 12 pM seeding molarity and sequenced on a HiSeq2500 in either Rapid Run Mode v2 or High Output Mode v3 in 2x100 paired-end format.

The raw sequencing reads were aligned to the ferret genome using star aligner (version 2.4.0g1). After read alignment, featureCounts (3) was used to quantify expression at the gene level. Genes with at least 5 reads in at least 4 samples were considered expressed and hence retained for further analysis. The gene level read counts data was normalized using the trimmed mean of M-values normalization (TMM) method (4) to adjust for sequencing library size differences. Normalized read counts were further adjusted for batch effect using a linear model. The residuals from the regression model were used for downstream analysis.

**Concentration dependence of IRE-1 inhibition in A549 cells.** A549 cells were seeded at a density of 120,000 cells in 35 mm glass-bottom dishes and grew in standard condition; 24 h later the cells were treated with IRE-1 inhibitor 4μ8C at 4, 8, 16, 32, 64, and 128 μM in the growth media for 24 h at 37°C. For control samples, the same amount of DMSO was added. The cells were then infected according to the infection assay described above and grown in infection media supplemented with the IRE-1 inhibitor 4μ8C (4, 8, 16, 32, 64, 128 μM, respectively) for 24 h before harvesting or fluorescence microscopy. For controls, the media was supplemented with the same amount of DMSO.

**XBP1 mRNA splicing assay.** RNA from the 4μ8C-treated and untreated A549 cells were extracted using miRNeasy mini kit (Qiagen) and reverse transcribed using the high capacity cDNA reverse transcription kit (ThermoFisher) according to the instructions. PCR amplification was performed using the XBP1 primers (forward: 5'-GGTAGATTTAGAAGAAGAGAACCAAAAACTTTTGCTAG-3'; reverse: 5'-TAAGGAACTGGGTCCTTCTGGG-3'), annealing temperature 69 °C, 35 cycles. The amplified DNA fragments were resolved on a 2.25% agarose gel in 0.5x TBE buffer with ethidium bromide.

*References:*

1. Hsu KL, Gildersleeve JC, & Mahal LK (2008) A simple strategy for the creation of a recombinant lectin microarray. *Mol Biosyst* 4(6):654-662.
2. Giles BM, *et al.* (2012) A computationally optimized hemagglutinin virus-like particle vaccine elicits broadly reactive antibodies that protect nonhuman primates from H5N1 infection. *J Infect Dis* 205(10):1562-1570.
3. Liao Y, Smyth GK, & Shi W (2014) featureCounts: an efficient general purpose program for assigning sequence reads to genomic features. *Bioinformatics* 30(7):923-930.
4. Robinson MD, McCarthy DJ, & Smyth GK (2010) edgeR: a Bioconductor package for differential expression analysis of digital gene expression data. *Bioinformatics* 26(1):139-140.

**Supplemental Figures:**

a

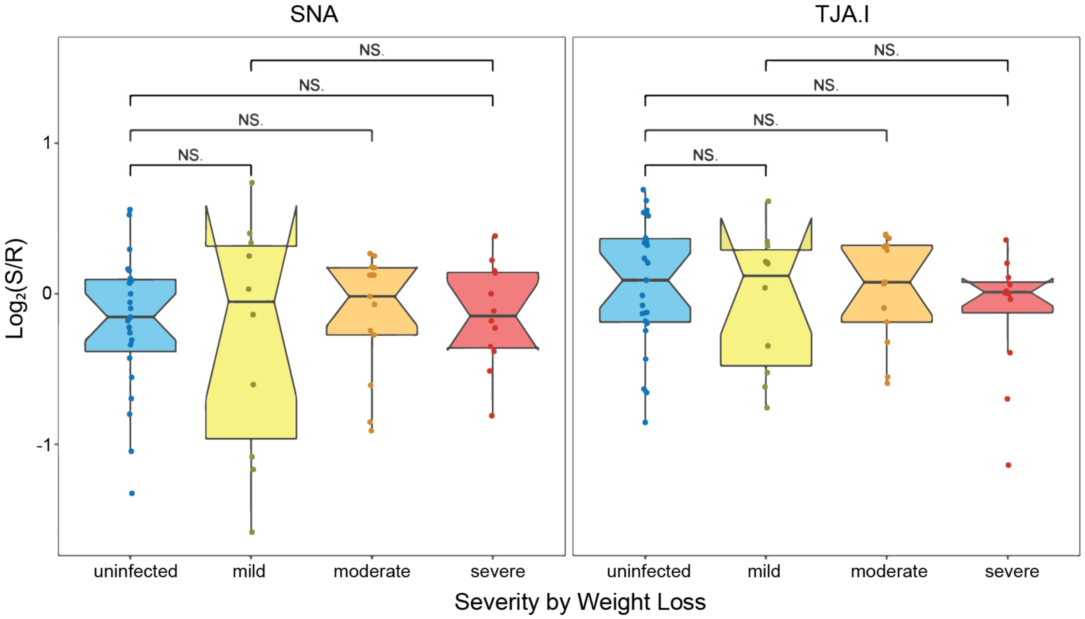


b

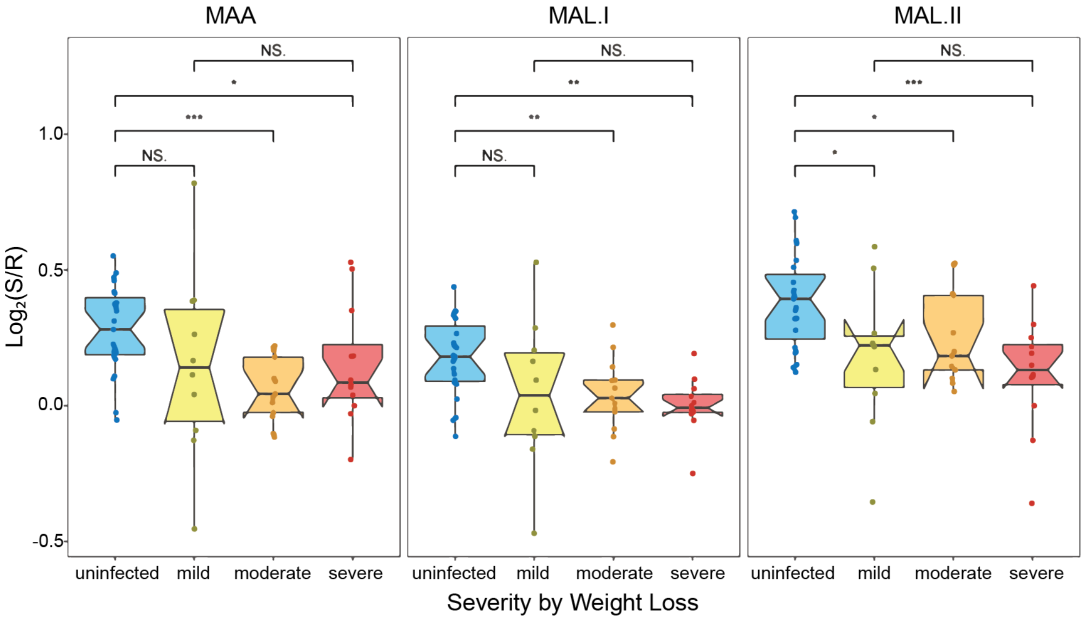


**Supplemental Figure 1: Impact of influenza infection on host sialosides.** a) Boxplots of binding by α-2,6-sialic acid lectins (SNA and TJA-I). Median normalized $\log_2$ ratios (Sample (S)/Reference (R)) of ferret lung samples were plotted. Ferret severity was indicated by colors (blue: uninfected, yellow: mild, orange: moderate, red: severe). N.S.: Not statistical, **: $p < 0.01$, ***: $p < 0.001$, Wilcoxon's t-test. b) Boxplots of binding by α-2,3-sialic acid lectins (MAA, MAL-I, MAL-II). Data shown is from ferrets 8 days post-infection.

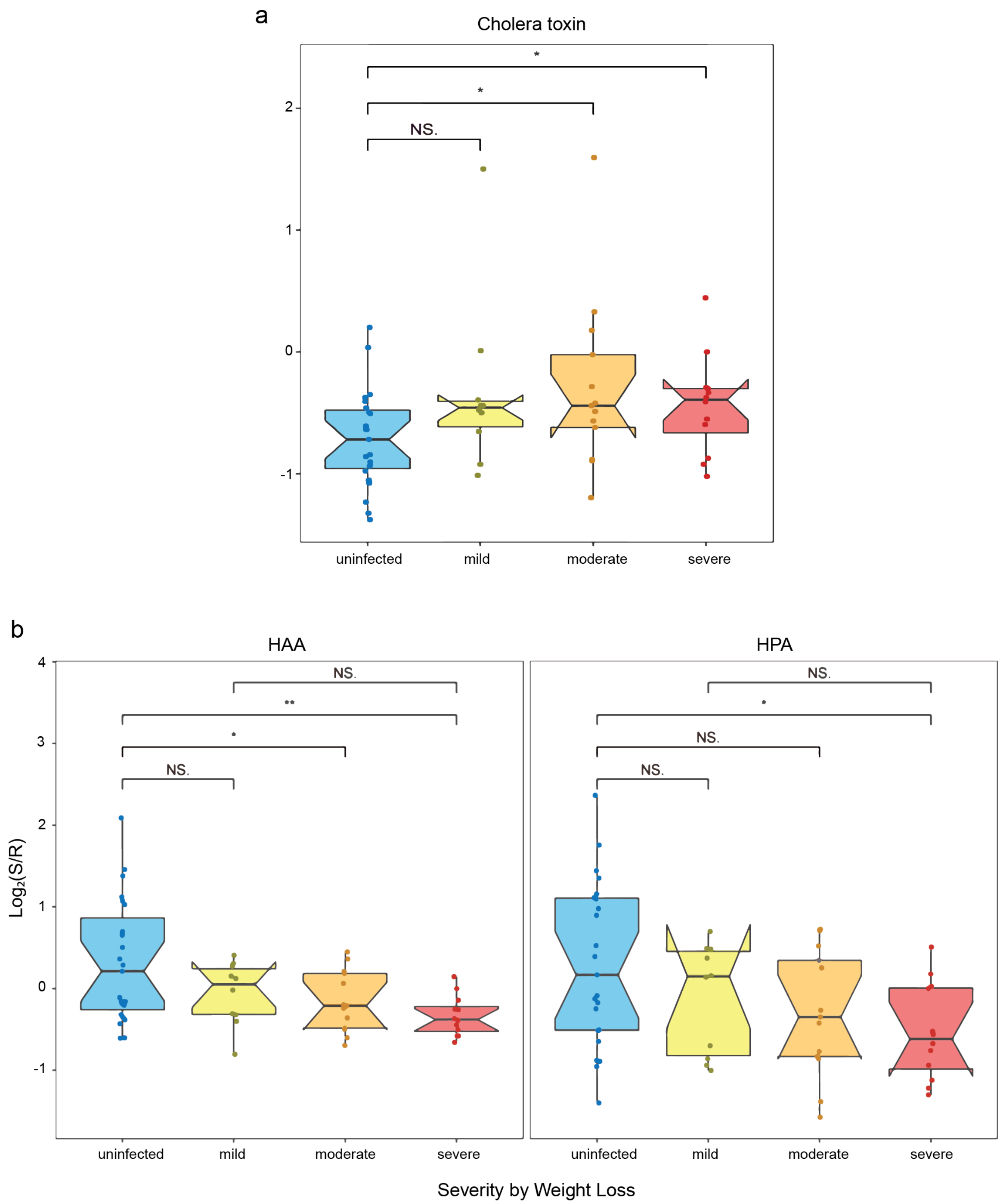


**Supplemental Figure 2: Impact of influenza infection on lipid raft marker GM1 and α-GalNAc.** a) Boxplot of GM1 change probed by cholera toxin. Median normalized $\log_2$ ratios (Sample (S)/Reference (R)) of ferret lung samples were plotted. b) Boxplots of α-GalNAc binding lectins (HAA, HPA). Ferret severity was indicated by colors (blue: uninfected, yellow: mild, orange: moderate, red: severe). N.S.: Not statistical, **: $p < 0.01$, ***: $p < 0.001$, Wilcoxon's t-test. Data shown is from ferrets 8 days post-infection.

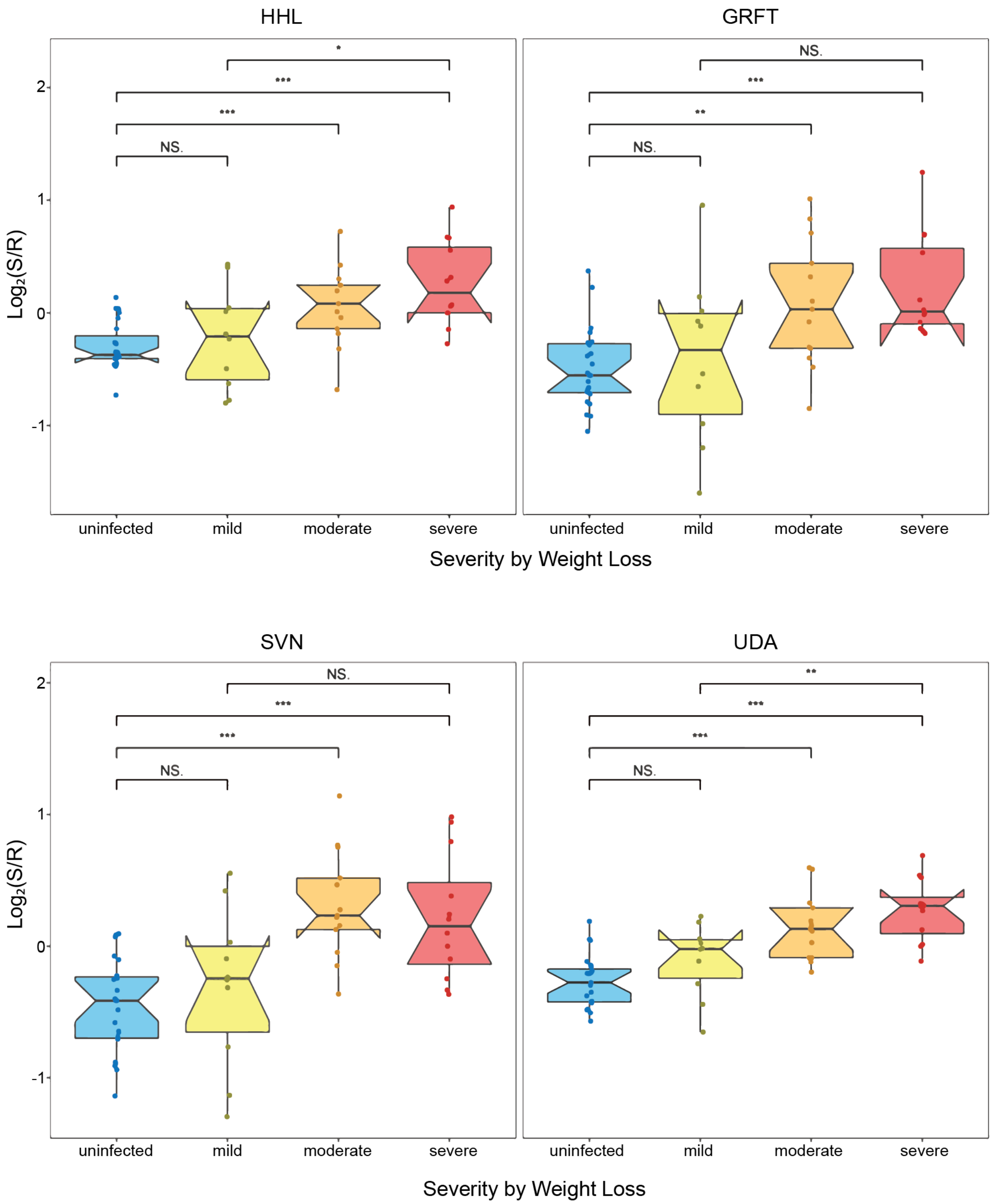


**Supplemental Figure 3: Impact of influenza infection on high mannose levels.** Boxplots of high mannose binding lectins (HHL, GRFT, SVN and UDA). Median normalized $log_2$ ratios (Sample (S)/Reference (R)) of ferret lung samples were plotted. Ferret severity was indicated by colors (blue: uninfected, yellow: mild, orange: moderate, red: severe). N.S.: Not statistical, **: $p < 0.01$, ***: $p < 0.001$, Wilcoxon's t-test. Data shown is from ferrets 8 days post-infection.

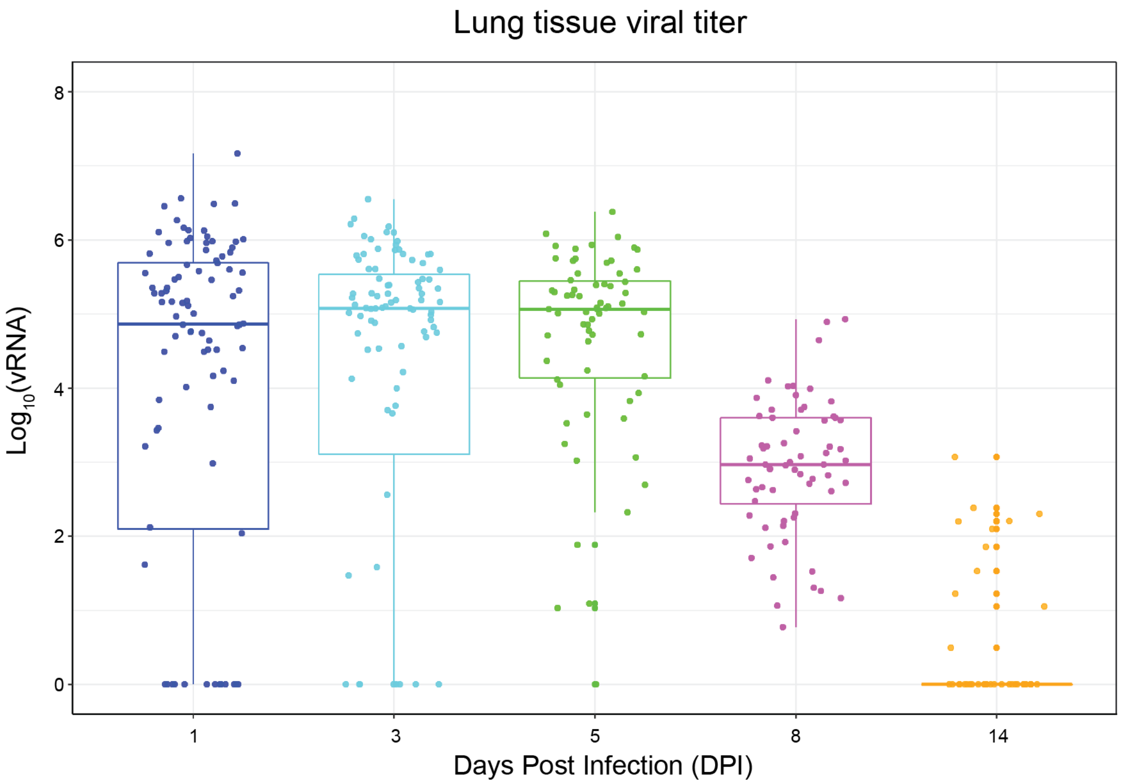


**Supplemental Figure 4: Boxplot of viral titer in ferret lung as a function of time.** $Log_{10}$ of viral titers from ferret lungs at t= 1, 3, 5, 8, and 14 days post infection.

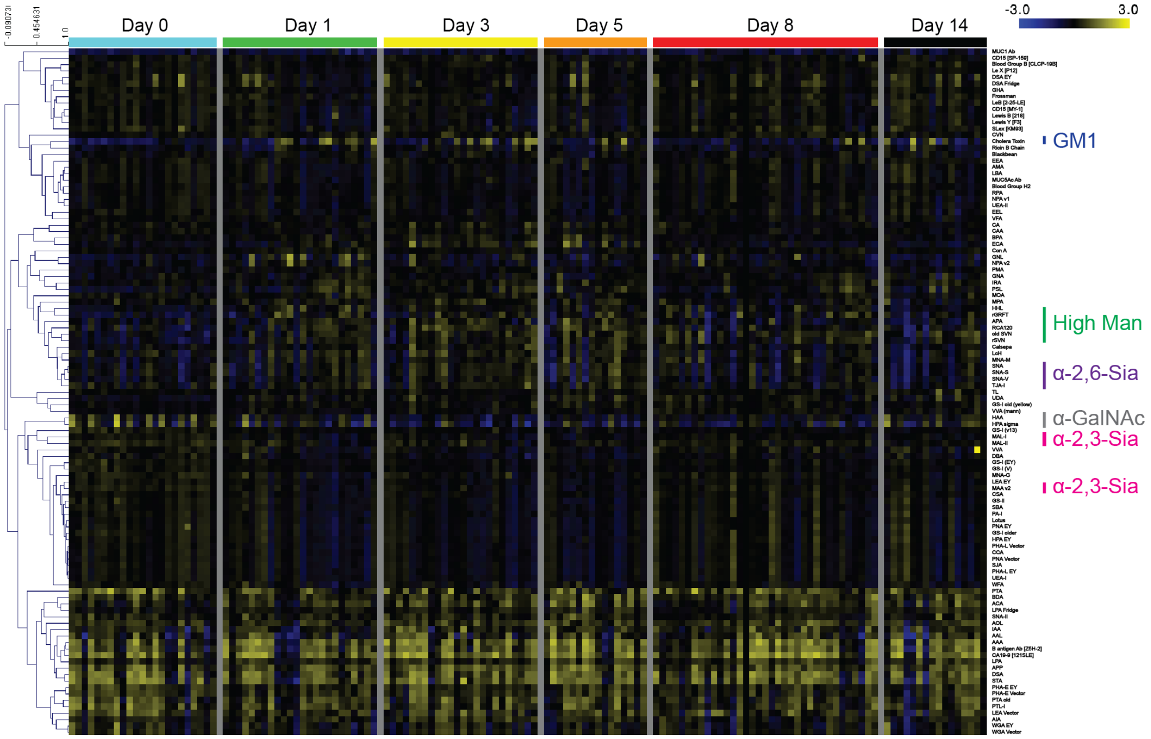


**Supplemental Figure 5: Heat map of lectin microarray data from all ferret samples** Median normalized $\log_2$ ratios (Sample (S)/Reference (R)) of ferret lung samples were organized by days post infection. (t=0, 1, 3, 5, 8, & 14 days; uninfected = day 0). Yellow, $\log_2(S) > \log_2(R)$; blue, $\log_2(R) > \log_2(S)$. Number of ferrets per: n=4, d0; n=12, d1-3; n=8, d 5; n=19, d 8; n=8, d 14). For uninfected (d0) ferrets 4 lung punch biopsies were used, for all other ferrets, 2 biopsies were analyzed. Lectins binding α-2,6-sialosides (purple), α-2,3-sialosides (pink), GM1 (blue), high mannose (green) and N-acetylgalactosamine (grey) are indicated on right.

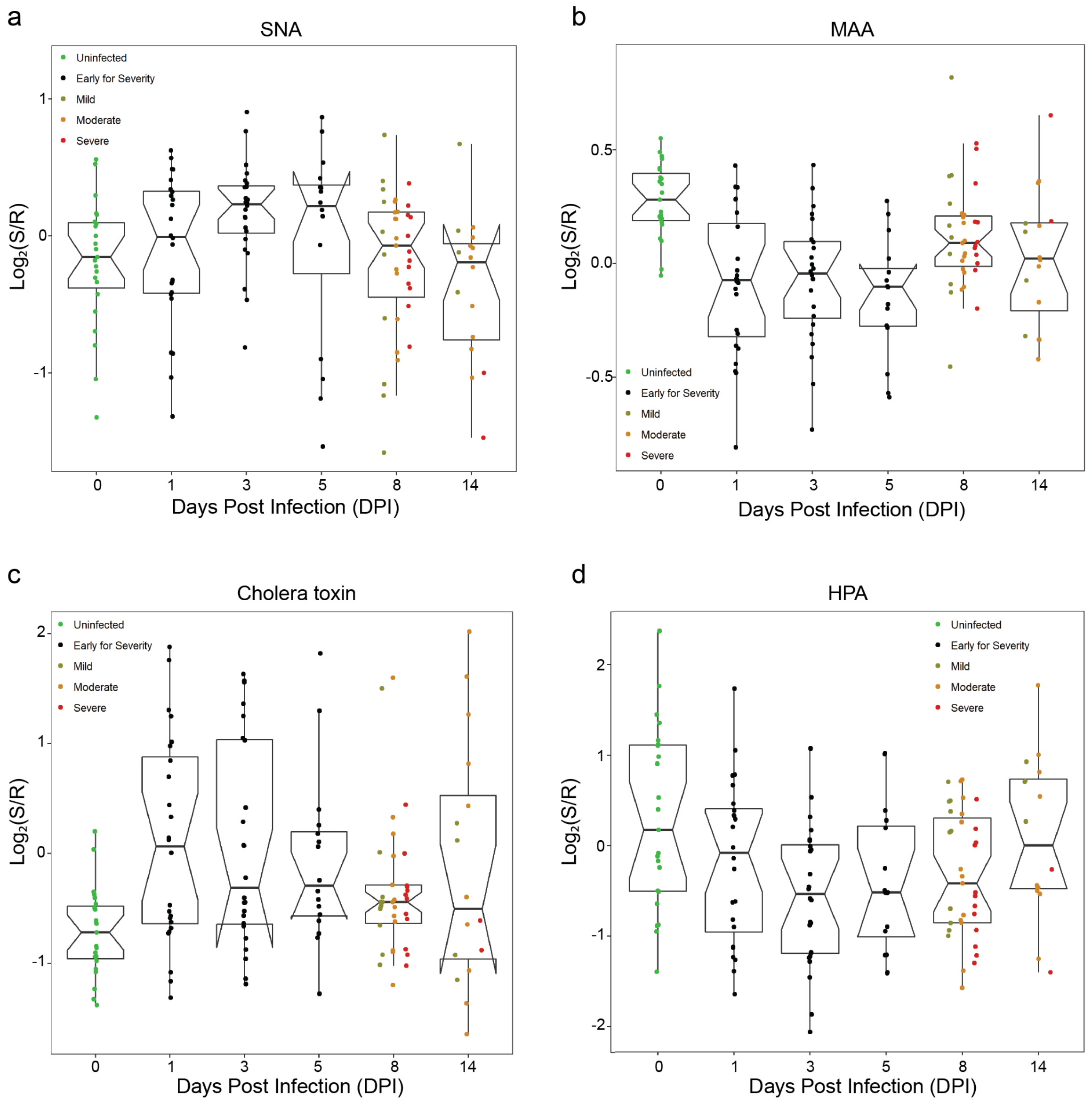


**Supplemental Figure 6: Host glycan dynamic change upon infection.** a) Boxplot of α-2,6-linked sialic acid change at different time points (0, 1, 3, 5, 8, 14 days post infection) probed by SNA. Median normalized $\log_2$ ratios (Sample (S)/Reference (R)) of ferret lung samples were plotted. Ferret severity is indicated by color (green: uninfected, black: early for severity, dark yellow: mild, orange: moderate, red: severe). b) Boxplot of α-2,3-linked sialic acid change at different time points (0, 1, 3, 5, 8, 14 days post infection) probed by MAA. c) Boxplot of GM1 change at different time points (0, 1, 3, 5, 8, 14 days post infection) probed by cholera toxin. d) Boxplot of α-GalNAc change at different time points (0, 1, 3, 5, 8, 14 days post infection) probed by HPA.

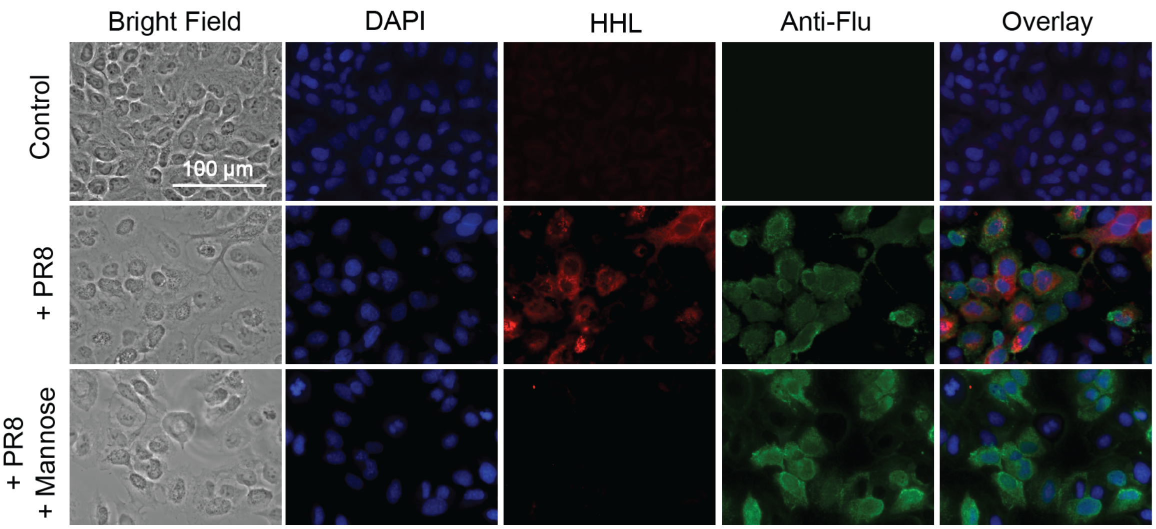


**Supplemental Figure 7: Fluorescence microscopy of A549 cells 24 h post infection.** Paraformaldehyde-fixed A549 cells were stained with biotinylated HHL developed with streptavidin CY5 (red), mouse anti-influenza nucleoprotein antibody developed with anti-mouse IgG CY3 (green), and DAPI (blue). High mannose inhibition was performed with incubation of the fixed cells with methyl mannose for 1 h at RT prior to infection. Bright field and overlay images of DAPI, HHL and anti-flu stained images were also shown. Scale bar: 100 µm. Three biological replicates were performed and at least six images were captured for each condition. These are representative pictures of the experiment results.

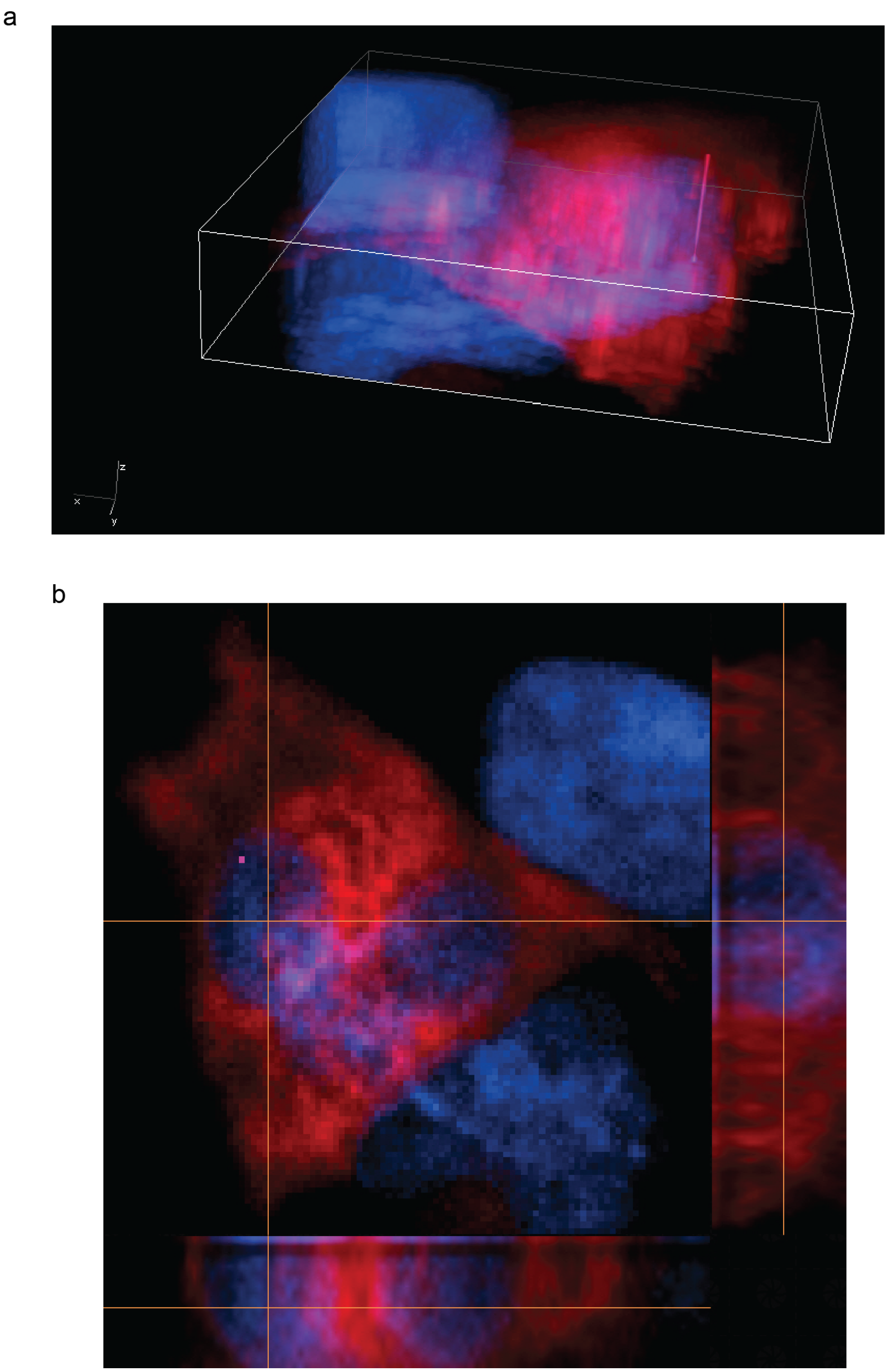

**Supplemental Figure 8: Fluorescence microscopy deconvolution image of the infected A549 cells.** Images were taken on fixed A549 cells 24 h post-infection by PR8. Blue: nucleus stain by DAPI, red: high mannose stain by HHL. a) Image in 3-D volume view. Width: 33.17 µm, height: 34.45 µm, depth: 6.98 µm. b) Image in XY, YZ, XZ slices view.

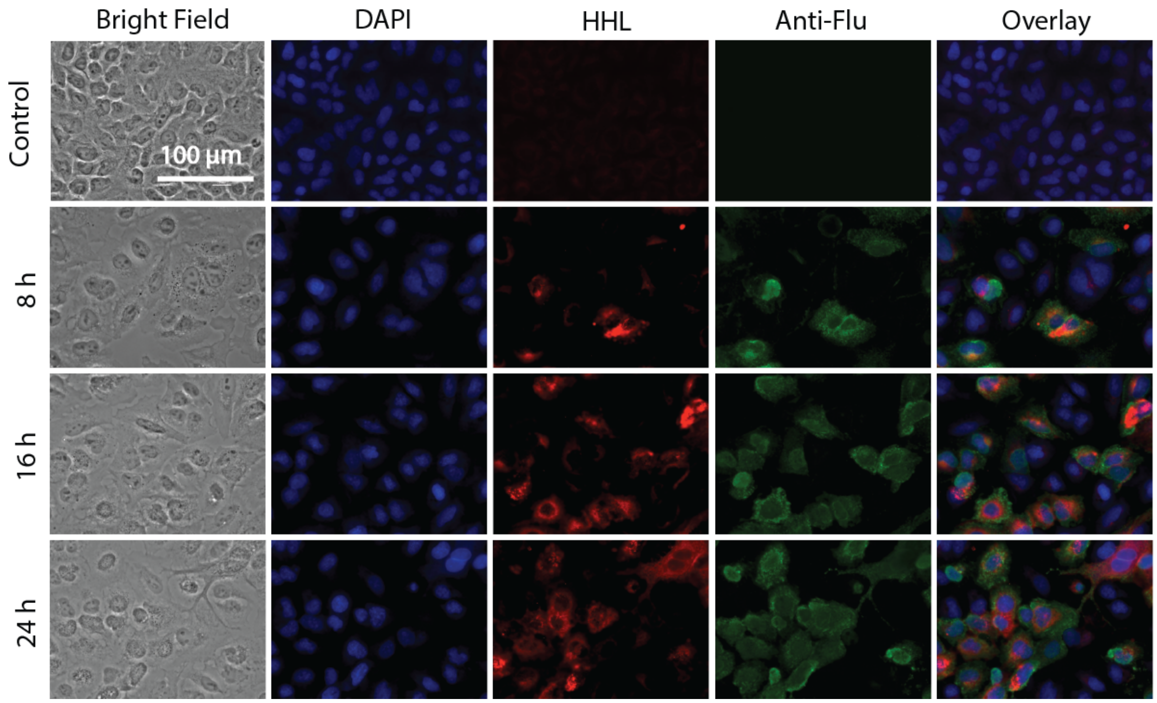


**Supplemental Figure 9: Fluorescence microscopy of A549 cells in the time course study on the influenza infection.** Cells grown for 8, 16, 24 h post infection were fixed for staining. Paraformaldehyde-fixed A549 cells were stained with biotinylated HHL developed with streptavidin Cy5 (red), mouse anti-influenza nucleoprotein (np) antibody developed with anti-mouse IgG Cy3 (green), and DAPI (blue). Bright field and overlay images of DAPI, HHL and anti-np stained images are also shown. Scale bar: 100 µm. Three biological replicates were performed and at least six images were captured for each condition. These are representative pictures of the experiment results.

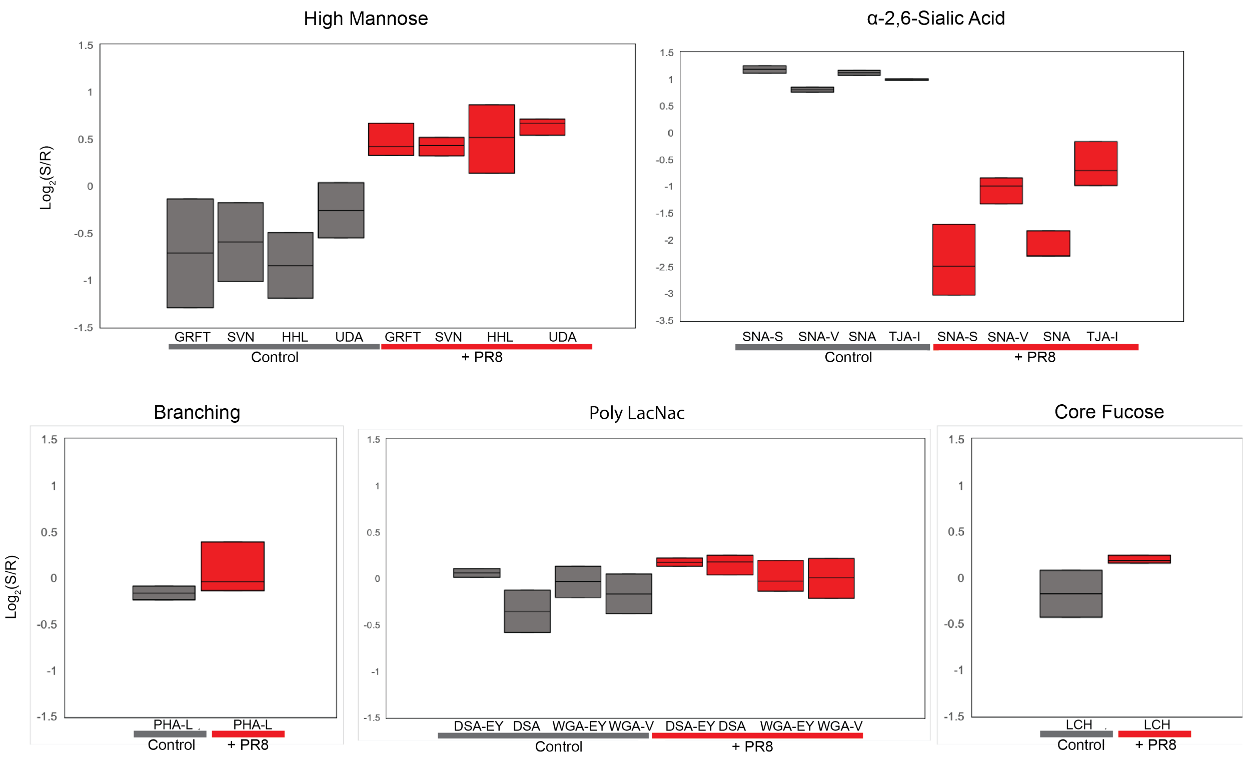


**Supplemental Figure 10: Graphical Analysis of Selectin Lectins from Fig. 5a.** Box and whisker plots of of high mannose binding (GRFT, SVN, HHL and and UDA), α-2,6-linked sialic acid binding (SNA from 3-sources, TJA-I), and complex binding (Branching: PHA-L, PolyLacNAc: DSA, WGA, Core Fucose: LcH) lectins. Median normalized $\log_2$ ratios (Sample (S)/Reference (R)) were plotted.

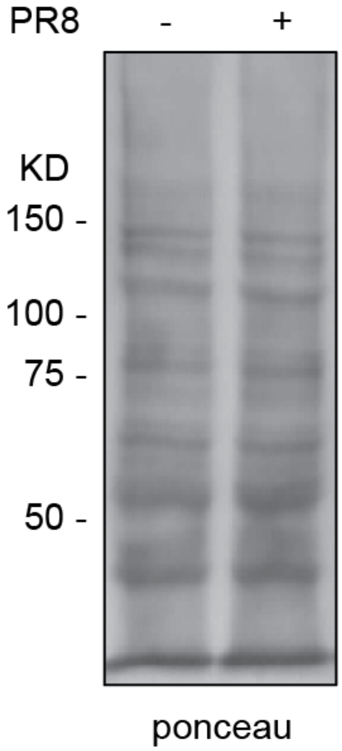


**Supplemental Figure 11: Ponceau staining of HHL lectin blot.** Duplicate lanes were run simultaneously and transferred to nitrocellulose. The blot was stained with ponceau solution at RT for 5 min followed by washing with deionized water for 3 times. Images were taken by photography using G:Box. The blot was then washed with PBST to remove ponceau for lectin staining.

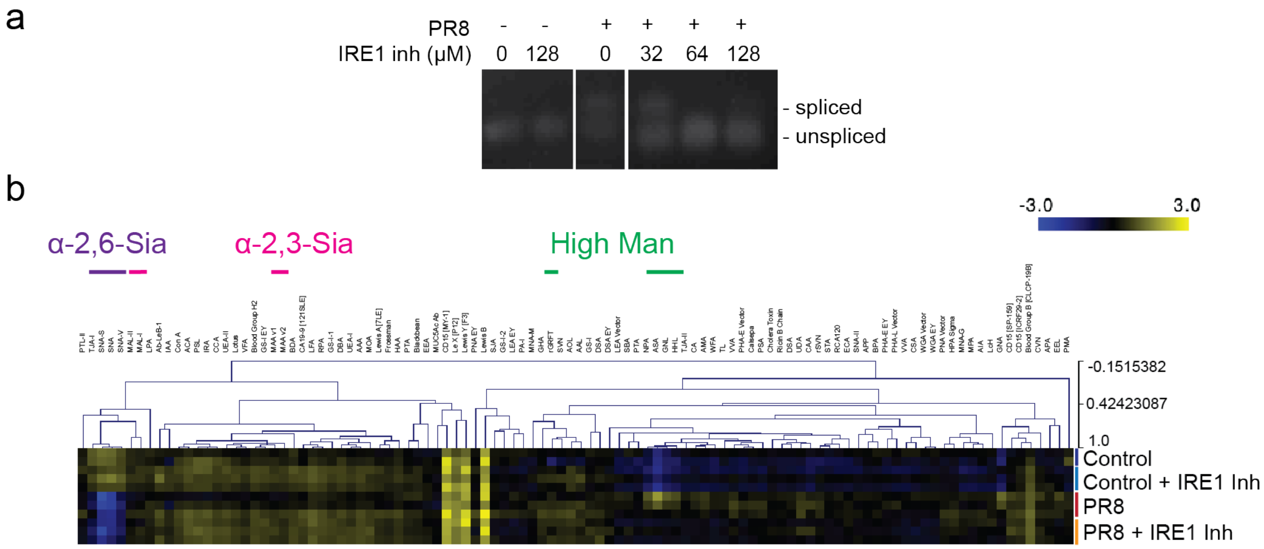


**Supplemental Figure 12: IRE1 inhibition.** a) Electrophoresis result of IRE1 inhibition condition test. A549 cells were treated with different concentrations (0, 32, 64, 128 μM) of IRE1 inhibitor 4μ8C 24 h prior to PR8 influenza virus infection. The spliced and unspliced forms are marked on the right. Three biological replicates were performed. A representative image is shown. b) Lectin microarray analysis of IRE-1 inhibition and influenza infection. Median normalized $\log_2$ ratios (Sample (S)/Reference (R)) of A549 cells were treated with IRE1 inhibitor 4μ8C (24 h, 64 μM, IRE1 Inh) prior to infection with PR8. Control cells were uninfected. Yellow, $\log_2(S) > \log_2(R)$; blue, $\log_2(R) > \log_2(S)$. Lectins binding α-2,6-sialosides (purple), α-2,3-sialosides (pink) and high mannose (green) are indicated on top.

**Supplemental Table 1: Lectin microarray print list.**

| Lectin | Species/Origin | Print Conc. (µg/mL) | Rough Specificity /Inhibitory monosaccharide | Vendor/ Source |
|---|---|---|---|---|
| **AAA** | *Anguilla anguilla* | 1000 | Fucose | EY |
| **AAL** | *Aleuria aurantia* | 500 | Fucose | Vector |
| **ACA** | *Amaranthus Caudatus* | 1000 | Gal-β1,3-GalNAc | Vector |
| **AIA** | *Artocarpus integrifolia* | 500 | β1,3-GalNAc | Vector |
| **AMA** | *Allium moly* | 500 | Oligo mannose | EY |
| **Anti-B.G. B [CLCP-19B]** | MAb mouse IgM [CLCP-19B] | undiluted | Blood Group B | Abcam |
| **Anti-B.G. H2** | MAb mouse IgM [A46-B/B10] | 200 | Blood Group H type 2 | Santa Cruz |
| **Anti-CA19-9 [121SLE]** | MAb mouse IgM [121SLE] | undiluted | Sialyl Lewis A | Abcam |
| **Anti-CD15 [ICRF29-2]** | MAb mouse IgM [ICRF29-2] | 500 | Lewis X | R and D Systems |
| **Anti-CD15 [MY-1]** | MAb mouse IgM [MY-1] | undiluted | Lewis X | Abcam |
| **Anti-Forssman** | MAb Rat IgM [M1/87.27.7HLK] | 200 | Forssman Antigen | Santa Cruz |
| **Anti-Le X [P12]** | MAb mouse IgM [P12] | undiluted | Lewis X | Abcam |
| **Anti-Le X [SP159]** | MAb rabbit IgG [sp159] | undiluted | Lewis X | Abcam |
| **Anti-Lewis B [218]** | IgM [T218] | undiluted | Lewis B | Abcam |
| **Anti-Lewis B [2-25-LE]** | MAb mouse IgG [2-25LE] | undiluted | Lewis B | Abcam |
| **Anti-Lewis Y [F3]** | MAb mouse IgM [F3] | undiluted | Lewis Y | Abcam |
| **Anti-SLeX [KM93]** | MAb mouse IgM [KM93] | undiluted | Sialyl Lewis X | CALBIOCHEM |
| **AOL** | *Aspergillus oryzae* | 1000 | Fucose | TCI |
| **APA** | *Abrus precatorius* | 500 | Gal-β1,3-GalNAc / Lac | EY |
| **APP** | *Aegopodium podagraria* | 1000 | GalNAc / Lac / Gal | EY |
| **ASA** | *Allium sativum* | 1000 | Mannose | Vector |
| **BDA** | *Bryonia dioica* | 1000 | GalNAc / Lac | EY |
| **Blackbean** | *Blackbean crude* | 1000 | GalNAc | EY |

| | | | | |
|---|---|---|---|---|
| **BPA** | *Bauhinia purpurea* | 500 | β-Gal / β-GalNAc | Vector |
| **CA** | *Colchicum autumnale* | 1000 | Bi-antennary N-link glycans | EY |
| **CAA** | *Caragana arborescens* | 500 | Bi-antennary N-link glycans | EY |
| **Calsepa** | *Calystegia sepium* | 500 | Bisecting N-link glycans | EY |
| **CCA** | *Cancer antennarius* | 1000 | 9-O-Acetly sialylation / 4-O-Acetyl sialylation | EY |
| **Cholera Toxin** | *Vibrio cholerae* | 2000 | Gal | Sigma |
| **Con A** | *Canavalia ensiformis* | 1000 | Tri-mannose core | Vector |
| **CSA** | *Cystisus scoparius* | 1000 | Terminal GalNAc | EY |
| **DBA** | *Dolichos Biflorus* | 500 | GalNAc | Vector |
| **diCBM40** | engineered nanI from *Clostridium perfringens* | 1000 | α Sialylation | Generated in house |
| **DSA** | *Datura stramonium* | 500 | LacNAc | EY/Vector |
| **ECA** | *Erythrina cristagalli* | 500 | LacNAc | Vector |
| **EEL/EEA** | *Eunonymus europaeus* | 1000 | Blood Group B | Vector/EY |
| **GafD** | recombinant GafD from *Escherichia coli* | 500 | GlcNAc | Generated in house |
| **GHA** | *Glechoma hederacea* | 500 | GalNAc | EY |
| **GNA/GNL** | *Galanthus nivalis* | 1000 | Oligo mannose | Vector/EY |
| **GS-I** | *Griffonia simplicifoia* | 1000 | α-Gal / Lac | Vector/EY |
| **GS-II** | *Griffonia simplicifoia-II* | 500 | GlcNAc | Vector |
| **HAA** | *Homarus americanus* | 1000 | Terminal GalNAc | EY |
| **HHL** | *Hippeastrum Hybrid* | 1000 | Oligo mannose | Vector |
| **HPA** | *Helix pomatia* | 500 | Blood Group A | EY/Sigma |
| **IAA** | *Iberis amara* | 500 | GalNAc | EY |
| **IRA** | *Iris Hybrid* | 1000 | GalNAc / Lac | EY |
| **LBA** | *Phaseolus lunatus* | 1000 | Blood Group A | EY |
| **LcH** | *Lens Culinaris* | 500 | Core Fucose | Vector |
| **LEA** | *Lycopersicon esculentum* | 500 | GlcNAc | Vector/EY |
| **LFA** | *Limax flavus* | 500 | α Sialylation | EY |
| **Lotus** | *Lotus tetragonolobus* | 500 | Fucose | Vector |

| | | | | |
|---|---|---|---|---|
| **LPA** | *Limulus polphemus* | 500 | α Sialylation | EY |
| **MAA** | *Maackia amurensis* | 500 | Sialylation/Sulfation | EY |
| **MAL-I** | *Maackia amurensis-I* | 1000 | Sialylation/Sulfation | Vector |
| **MAL-II** | *Maackia amurensis-II* | 1000 | Sialylation/Sulfation | Vector |
| **MNA-G** | *Morus nigra Morniga G* | 500 | GalNAc | EY |
| **MNA-M** | *Morus nigra Morniga M* | 500 | Oligo mannose | EY |
| **MOA** | *Marasmium oreades* | 500 | Gal-α1,3 | EY |
| **MPA** | *Maclura pomifera* | 500 | β1,3-GalNAc | Vector |
| **NPA** | *Narcissus pseudonarcissus* | 500 | Oligo mannose | Vector |
| **PA-I** | *Pseudomonas aeruginosa* | 500 | Gal | Sigma |
| **PHA-E** | *Phaseolus vulgaris Erythroagglutinin* | 500 | Complex N-link glycans | Vector/EY |
| **PHA-L** | *Phaseolus vulgaris Leukoagglutinin* | 500 | β1,6 Branching N-Link glycans | Vector/EY |
| **PMA** | *Polygonatum multiflorum* | 500 | Oligo mannose | EY |
| **PNA** | *Arachis hyogaea* | 500 | Gal-β1,3-GalNAc | Vector/EY |
| **PSA** | *Pisum sativum* | 500 | Core Fucose | Vector |
| **PSL** | *Polyporus squamosus* | 1000 | α2,6 sialylation | EY |
| **PTA** | *Psophocarpus tetragonolobus* | 500 | Blood Groups | EY |
| **PTL-I** | *Psophocarpus tetragonolobus-I* | 1000 | Blood Group A | Vector |
| **PTL-II** | *Psophocarpus tetragonolobus-II* | 1000 | α2 Fucose | Vector |
| **PWM** | *Phytolacca americana* | 1000 | GlcNAc | EY |
| **RCA120** | *Ricinus Communis Agglutinin I* | 1000 | Gal / Lac | Vector |
| **rCVN** | *recombinant Cyanovirin* | 1000 | High mannose | Gift from O'Keefe |
| **rGRFT** | *recombinant Griffithsin* | 1000 | High mannose | Gift from O'Keefe |
| **Ricin B Chain** | *Ricinus communis* | 1000 | Gal | Vector |

| | | | | |
|---|---|---|---|---|
| **RPA** | *Robinia pseudoacacia* | 500 | Complex N-link glycans | EY |
| **rSVN** | *recombinant Scytovirin* | 500 | High mannose | Gift from O'Keefe |
| **SBA** | *Glycine max* | 500 | LacdiNAc | Vector |
| **SJA** | *Sophora japonica* | 1000 | LacdiNAc | Vector |
| **SNA** | *Sambucus nigra* | 500 | α2,6 sialylation | Vector/EY/ Sigma |
| **SNA-II** | *Sambucus nigra-II* | 500 | α2 Fucose /oligo mannose | EY |
| **STA** | *Solanus tuberosum* | 500 | GlcNAc | Vector |
| **TJA-I** | *Trichosanthes japonica-I* | 1000 | α2,6 sialylation | TCI |
| **TJA-II** | *Trichosanthes japonica-II* | 500 | α2 Fucose | TCI |
| **TL** | *Tulipa sp.* | 1000 | GlcNAc | EY |
| **UDA** | *Urtica dioica* | 1000 | GlcNAc / Oligo mannose | EY |
| **UEA-I** | *Ulex europaaeus-I* | 1000 | α2 Fucose | Vector |
| **UEA-II** | *Ulex europaaeus-II* | 1000 | GlcNAc | Vector |
| **VFA** | *Vicia faba* | 1000 | GlcNAc | EY |
| **VVA** | *Vicia villosa* | 500 | Terminal GalNAc | Vector/EY |
| **WFA** | *Wisteria floribunda* | 500 | GalNAc-β1,4 | Vector |
| **WGA** | *Triticum vulgare* | 1000 | GlcNAc | Vector/EY |
